# Programmable low-cost DNA-based platform for viral RNA detection

**DOI:** 10.1101/2020.01.12.902452

**Authors:** Lifeng Zhou, Arun Richard Chandrasekaran, Jibin Abraham Punnoose, Gaston Bonenfant, Stephon Charles, Oksana Levchenko, Pheonah Badu, Cassandra Cavaliere, Cara T. Pager, Ken Halvorsen

## Abstract

Viral detection is critical for controlling disease spread and progression. Recent emerging viral threats including Zika, Ebola, and the current COVID-19 outbreak highlight the cost and difficulty in responding rapidly. To address these challenges, we develop a platform for low-cost and rapid detection of viral RNA with DNA nanoswitches designed to mechanically reconfigure in response to specific viruses. Using Zika virus as a model system, we show non-enzymatic detection of viral RNA to the attomole level, with selective and multiplexed detection between related viruses and viral strains. For clinical-level sensitivity in biological fluids, we paired the assay with a sample preparation step using either RNA extraction or isothermal pre-amplification. Our assay can be performed with minimal or no lab infrastructure, and is readily adaptable to detect other viruses. We demonstrate the adaptability of our method by quickly developing and testing DNA nanoswitches for detecting a fragment of SARS-CoV-2 RNA in human saliva. Given this versatility, we expect that further development and field implementation will improve our ability to detect emergent viral threats and ultimately limit their impact.

---

Newly emerging or re-emerging viruses pose significant challenges to health care systems, particularly as globalization has contributed to the rampant spread of these viruses.^1^ RNA viruses are frequently the cause of sweeping outbreaks as these viruses have high mutation rates and thus evolve rapidly.^2,3^ Examples of this include the annual influenza outbreak, Ebola virus, Zika virus (ZIKV) and the SARS-CoV-2 virus responsible for the COVID-19 pandemic. Technological advancements in structural biology and genomics have been important for identifying viruses, and for advancing fundamental viral research and antiviral therapeutics.^4^ However, clinical methods for robust, low-cost and rapid detection of viral infections remain a major challenge for emergent viruses, especially in resource limited areas.

Detection of RNA viruses in the clinical setting is typically performed using either immunological detection based on enzyme-linked immunosorbent assay (ELISA) to detect IgM antibodies or nucleic acid testing (NAT) based on a reverse transcription polymerase chain reaction (RT-PCR) assay to detect viral RNA.^5–8^ Diagnosing RNA viruses is made challenging by several factors including a limited time window for detection, low or varying viral load, cross-reactive IgM antibodies, and laboratory resources. The detection time windows can vary widely from as short as several days to as long as several months,^5^ and molecular detection techniques are usually most reliable if performed within the first two weeks of the disease.^9,10^ Depending on the timing of testing relative to infection, even highly sensitive NAT assays may still produce false negative or positive results.^6,11^ On the other hand, results from IgM serology tests often cannot distinguish related viruses or different strains of the same virus due to cross-reactivity of IgM antibodies, thus leading to false positive results.^12,13^ These detection challenges are further exacerbated when outbreaks occur in low resource settings where infrastructure for these lab-intensive tests can be lacking, accelerating the spread of disease.^7,14,15^

In response to some of these challenges, new techniques are being developed to detect emerging viruses. Among these are methods that adopt nanoparticles,^16,17^ graphene-based biosensors,^18,19^ and CRISPR-based methods,^20-22^ to name a few. Many of these proposed strategies, although based on cutting-edge technology, require multiple reactions or signal transformation steps. Here, we addressed these biosensing challenges by developing an assay that uses programmable DNA nanoswitches^23,24^ for detection of viral RNA at clinically relevant levels. We validate our strategy using ZIKV as a proof-of-concept system given its high global health relevance and continued threat due to its re-emerging mosquito-borne nature. Although ZIKV is typically associated with mild symptoms, it has been linked to devastating birth defects associated with intrauterine infections, appearance of Guillian-Barré syndrome in adults, and the possibility of sexual transmission.^7,10^ Moreover, despite significant advances in understanding the molecular biology of ZIKV, there is still a lack of antiviral drugs and vaccines, making robust detection of ZIKV vital to controlling the spread of the disease and implementing early treatments.^25^

Our strategy for detecting the presence of viral RNA is based on using DNA nanoswitches that have been designed to undergo a conformational change (from linear to looped) upon binding a target viral RNA sequence (**Fig. 1a**). The presence of the viral RNA would be indicated by shifted migration of the looped nanoswitch by gel electrophoresis. Importantly, the system is designed to use common nucleic acid staining of the nanoswitch itself that can intercalate thousands of dye molecules to provide an inherently strong signal. Previously, we demonstrated sensitive and specific detection of DNA oligonucleotides^26^ and microRNAs (~22 nucleotides long)^27^ using this approach. Applied here to viral RNA detection, we solved the challenges of detecting a long viral RNA (>10,000 nucleotides), improving the signal for long RNAs with a new signal multiplication strategy, and developing workflows for measuring viral loads in biological and mock clinical samples. We showed how multiplexing can be used to detect multiple viruses simultaneously from a single sample and demonstrated high specificity even between closely-related strains of Zika. In addition, as a quick response to the COVID-19 pandemic, we developed DNA nanoswitches in two days (**Fig. 1b**) and demonstrated the detection of SARS-CoV-2 RNA in human saliva at clinically relevant concentrations. Our approach is non-enzymatic, but can optionally be combined with an isothermal amplification step, allowing use in low resource areas (**Fig. 1c**). This work enables direct detection of viral RNA without amplification and paves the way toward a low-cost assay for detection of RNA viruses.

**Fig. 1.**
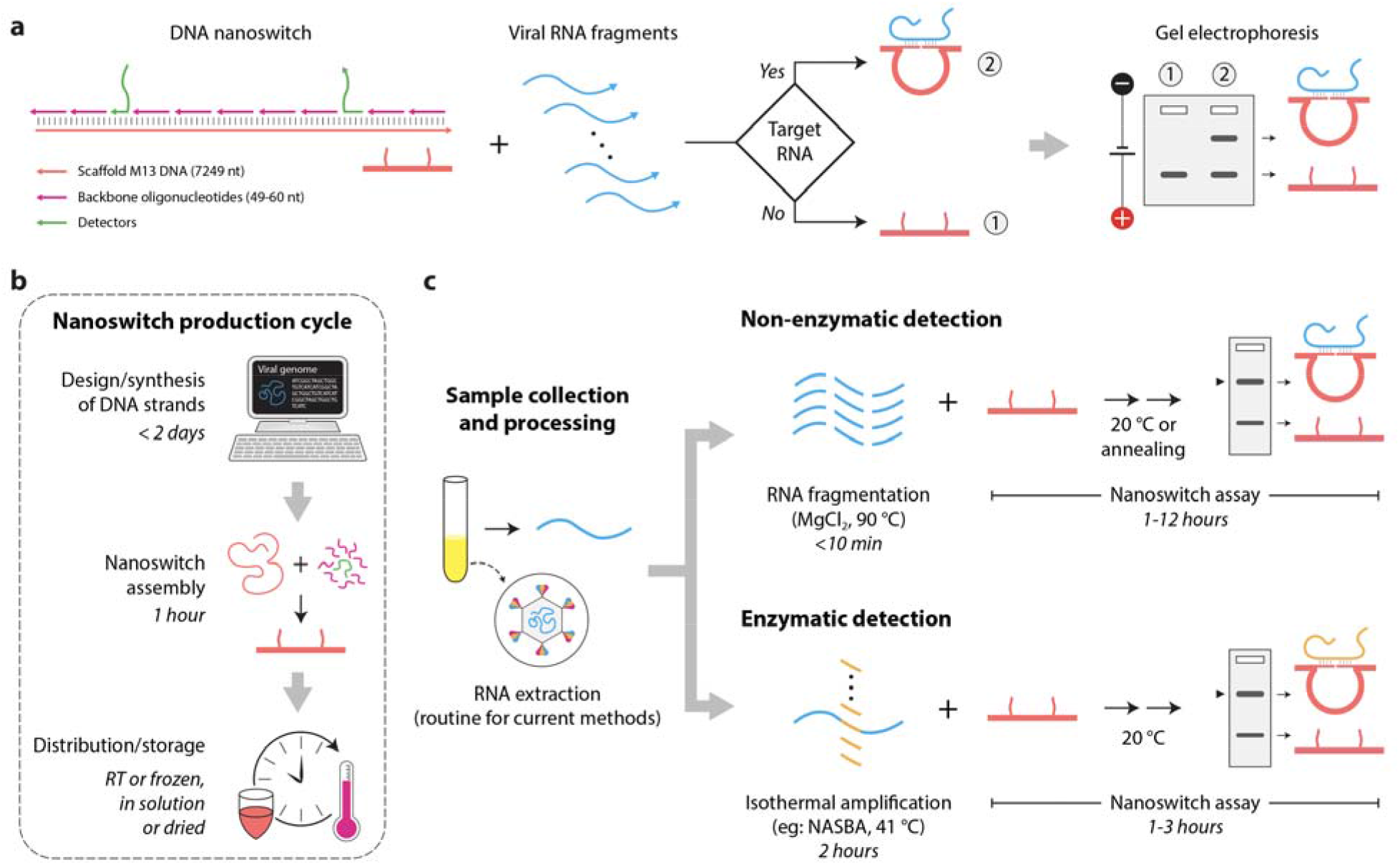
Detection of viral RNA using DNA nanoswitches. **(a)** Schematic of the DNA nanoswitch and detection of a viral RNA sequence. **(b)** Fast development cycle of nanoswitches for RNA viruses. **(c)** Nanoswitch based assay allows direct detection using a non-enzymatic approach (top panel) and can optionally be combined with an isothermal amplification step (bottom panel).

## Results

As a first proof-of-concept for detecting ZIKV, we designed DNA nanoswitches to target an already validated sequence in the ZIKV genome that has been used to bind primers in qPCR^28^ (all oligo sequences are specified in **Tables S1 to S10**). We made the DNA nanoswitches by hybridizing single-stranded DNA (ssDNA) oligos to linearized single-stranded M13mp18 (M13) genomic DNA in a thermal annealing ramp for 1 hour^24^ and purified them by high-performance liquid chromatography (HPLC).^29^ For our initial detection target, we *in vitro* transcribed RNA from the pFLZIKV infectious plasmid containing the full length genome of the Cambodia ZIKV isolate (FSS13025) (**Fig. S1)**.^30^ Previous results have shown robust nanoswitch detection of small DNA and RNA sequences (20-30 nucleotides), but the long viral RNA is expected to have strong secondary structures that may interfere with our detection.^31,32^ To overcome this, we used a chemical fragmentation method to segment the RNA into small pieces that are mostly shorter than 200 nt (**Fig. 2a-b** and **Fig. S2**). By incubating with our nanoswitch in an annealing temperature ramp, we showed successful detection of the fragmented viral RNAs by gel electrophoresis, thus validating our approach (**Fig. 2c**).

**Fig. 2.**
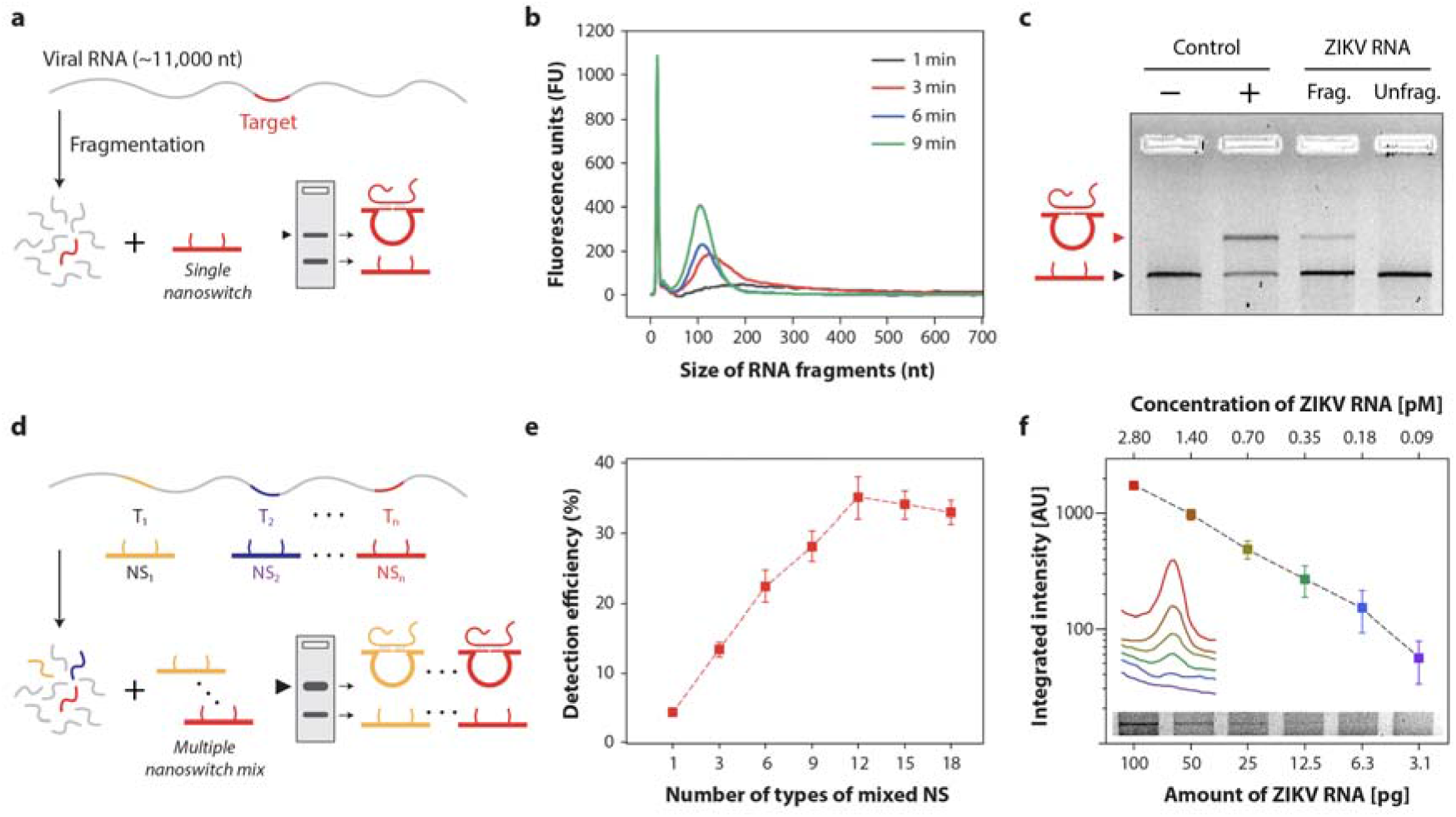
Detection of viral RNA using DNA nanoswitches. (**a)** Schematic of the fragmentation of viral RNA and subsequent detection by the DNA nanoswitch. **(b)** Fragmentation analysis of ZIKV RNA that was fragmented at 94 °C for 1, 3, 6, and 9 minutes. (**c)** Proof-of-concept showing detection of a target region chosen from the literature.^28^ (**d)** Schematic of the design of multiple nanoswitches for detection with the signal multiplication strategy. (**e)** Validation of the signal multiplication strategy: the detection signal was increased for a fixed pool of DNA targets when using multiple targeting nanoswitches. (**f)** Detection sensitivity of the pooled nanoswitches for ZIKV RNA. Error bars represent standard deviation from triplicate experiments.

Having shown successful detection of ZIKV RNA using a single target sequence, we recognized that we could exploit the large genome size (~ 11,000 nucleotides) to increase our detection signal through multiple targets. Once the long viral RNA is fragmented, the number of available target sequences increases dramatically. Since our detection signal is proportional to the number of looped nanoswitches, a nanoswitch mixture for different target sequences within the viral genome is expected to provide an increased signal. To test this, we developed an algorithm for choosing multiple sequence regions in the viral genome that can be targeted by the nanoswitches (**Note S1**). First, we chose the default target length as 30 nucleotides based on results from screening nanoswitches with different detection arm lengths (**Fig. S3**). Then, the algorithm selectively excluded target sequences that could form stable secondary structures (**Fig. S4**) and cross-binding with nanoswitch backbone oligos (**Fig. S5**), and enforced GC content and uniqueness of sequences. Based on these criteria, we chose 18 target regions along the entire ZIKV RNA for testing and designed the nanoswitches. To facilitate use of our Matlab-based software, we have built a graphical user interface (**Fig. S6**) and made it freely available (**File S1**).

We then validated quality and function of each nanoswitch in the panel of 18 nanoswitches. All nanoswitches performed well with a molar excess of positive DNA controls, although they showed more signal variation with fragmented ZIKV RNA (**Fig. S7**). We ranked the nanoswitches from strongest to weakest signal and made a series of equimolar nanoswitch mixtures. Using these mixtures, we validated our inherent signal multiplication strategy using a low concentration pool of equimolar DNA fragments to mimic the fragmented RNA. We observed that our detection signal increased steadily up to around 12 nanoswitches (**Fig. 2d-e**), and then plateaued above that value. This plateau was not unexpected considering that the largest mixtures added lower performing nanoswitches that may contribute less to the overall sample. Since there was no significant change in performance between 12 and 18, we continued using the 18 nanoswitches mix for our follow up experiments.

High sensitivity is one of the key requirements for viral detection. Clinical levels of ZIKV RNA in body fluids of infected patients are often in the femtomolar range,^7,21,33^ making amplification a prerequisite for most detection approaches. Based on our earlier observation that DNA nanoswitches can detect microRNAs (~22 nucleotides) in the sub-picomolar scale^27^ without amplification, we wanted to assess the sensitivity of our approach for ZIKV RNA detection. We reacted the DNA nanoswitch mixture with different amounts of fragmented RNA in a 12-hour annealing temperature ramp from 40 °C to 25 °C. The results showed visible detection for ZIKV RNA as low as 12.5 pg (~3.5 attomole) (**Fig. 2f** and **Fig. S8**) in a 10 μl reaction volume. Consistent with **Fig. 2e**, the approach based on using a nanoswitch mix outperformed the highest performing nanoswitch used as a single agent, which had visible detection to about 50 pg (~14 attomole) (**Fig. S9**).

Another key requirement for a clinical viral detection assay is specificity. Since ZIKV and Dengue virus (DENV) have overlapping geographical distributions and clinical symptoms, infection with either virus may result in clinical misdiagnosis.^34^ Serological diagnostic assays are known to show antibody crossreactivity between the two viruses, and DENV has some similarity to ZIKV in its envelope protein^12^ and genome sequence.^21,31^ To test the specificity of our approach, we designed a similar panel of nanoswitches to detect DENV (**Fig. S10**). Using the pooled nanoswitches specific for ZIKV and DENV, we mixed each set with *in vitro* transcribed RNA from each virus and found perfect specificity, with each assay only detecting its correct target RNA **(Fig. 3a).** Using the programmability of the nanoswitch, we further demonstrated a multiplexed system for simultaneous detection of ZIKV and DENV. In this case we modified the DENV responsive nanoswitches to form a smaller loop size (**Fig. S11**), causing two distinct detection bands to migrate to different positions in the gel. Specifically, ZIKV RNA-nanoswitch complex migrated slower/higher in the gel, while the complex of DENV RNA and the nanoswitch migrated faster/lower in the gel (**Fig. 3b**). Therefore, in a single reaction our nanoswitch showed differential and specific detection of ZIKV and DENV RNA. By programming different loop sizes for different targets, this assay can be expanded for up to five viral targets.^27^

**Fig. 3.**
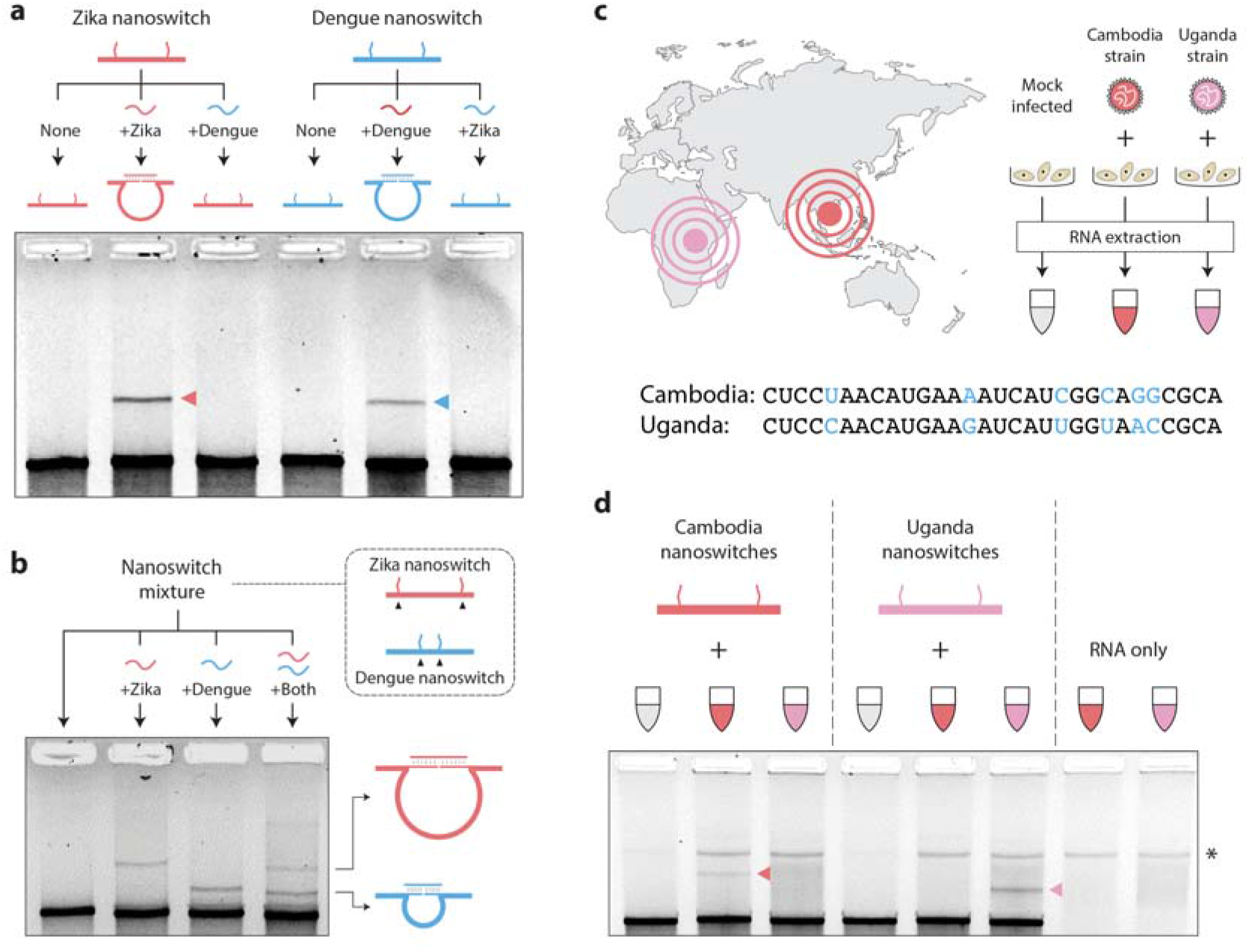
DNA nanoswitches specifically and differentially detect RNA from two different flaviviruses and between two highly similar ZIKV isolates. **(a)** ZIKV nanoswitches specifically detect ZIKV RNA but not DENV RNA, and *vice versa*. (**b)** Multiplexed detection of ZIKV and DENV RNA. (**c)** Illustration showing culture a**nd** RNA extraction of ZIKV Cambodia and Uganda strains. The mismatches in a representative target sequen**ce** between the two strains are shown. (**d)** Specificity test of Cambodia and Uganda strains of ZIKV RNA. * denotes a band of contaminating cellular DNA following RNA isolation.

In addition to possible misdiagnosis between different viruses, there is an additional challenge in determining the specific strain of a virus. For example, in Latin America four different DENV serotypes are known to be present and co-circulate, where misdiagnosis of the infecting strain can have significant implications for treatment options.^35^ Thus being able to accurately identify a circulating strain of virus broadly impacts medical care, surveillance and vector control.^36^ ZIKV was first identified in Uganda in 1947 before spreading to Asia and the Americas, and ZIKV strains (classified within African or Asian lineage) share significant sequence homology.^37^ To investigate if our assay can distinguish between the Asian and African lineages, we tested our nanoswitches against two ZIKV strains which have an ~89% sequence homology, namely the FSS13025 isolated from Cambodia and the MR766 strain isolated from Uganda. In designing the ZIKV strain-specific nanoswitches, we identified five target regions that each have a 5-6 nucleotide difference (**Fig. 3c** and **Fig. S12**). To achieve better discernment of the detection signal, the nanoswitches for the Uganda strain were designed to form a smaller loop-size than those designed for Cambodia. Next, a human hepatocellular carcinoma cell line (Huh7) was infected with either the Cambodian or Ugandan ZIKV strain. Infected cells were processed to extract total RNA, which was then fragmented and incubated with nanoswitches to probe for viral RNA from either the ZIKV Cambodia or ZIKV Uganda infected cells. The results showed that our assay was able to discriminate between two strains of the same virus even with high genetic similarities (**Fig. 3d** and **Fig. S12**).

Further applying our technique to detect ZIKV RNA in biological samples, we either mock-infected or infected Huh7 cells with the Cambodia ZIKV strain at a multiplicity of infection of 1 and extracted RNA from the ZIKV infected cells at 1-, 2- and 3-days post-infection.^38^ The nanoswitch assay detected ZIKV viral RNA from the infected cells but not the mock infected cells (**Figs. 4a-b** and **Fig. S13**). Our detection result shows that the copies of ZIKV RNA within infected cells steadily increased upon the infection and plateaued at 2- and 3-days post-infection (**Fig. 4c**). These data demonstrate that our assay can detect ZIKV RNA in infected cell lines, and in contrast to typical RT-PCR assays without amplification of the viral RNA.

**Fig. 4.**
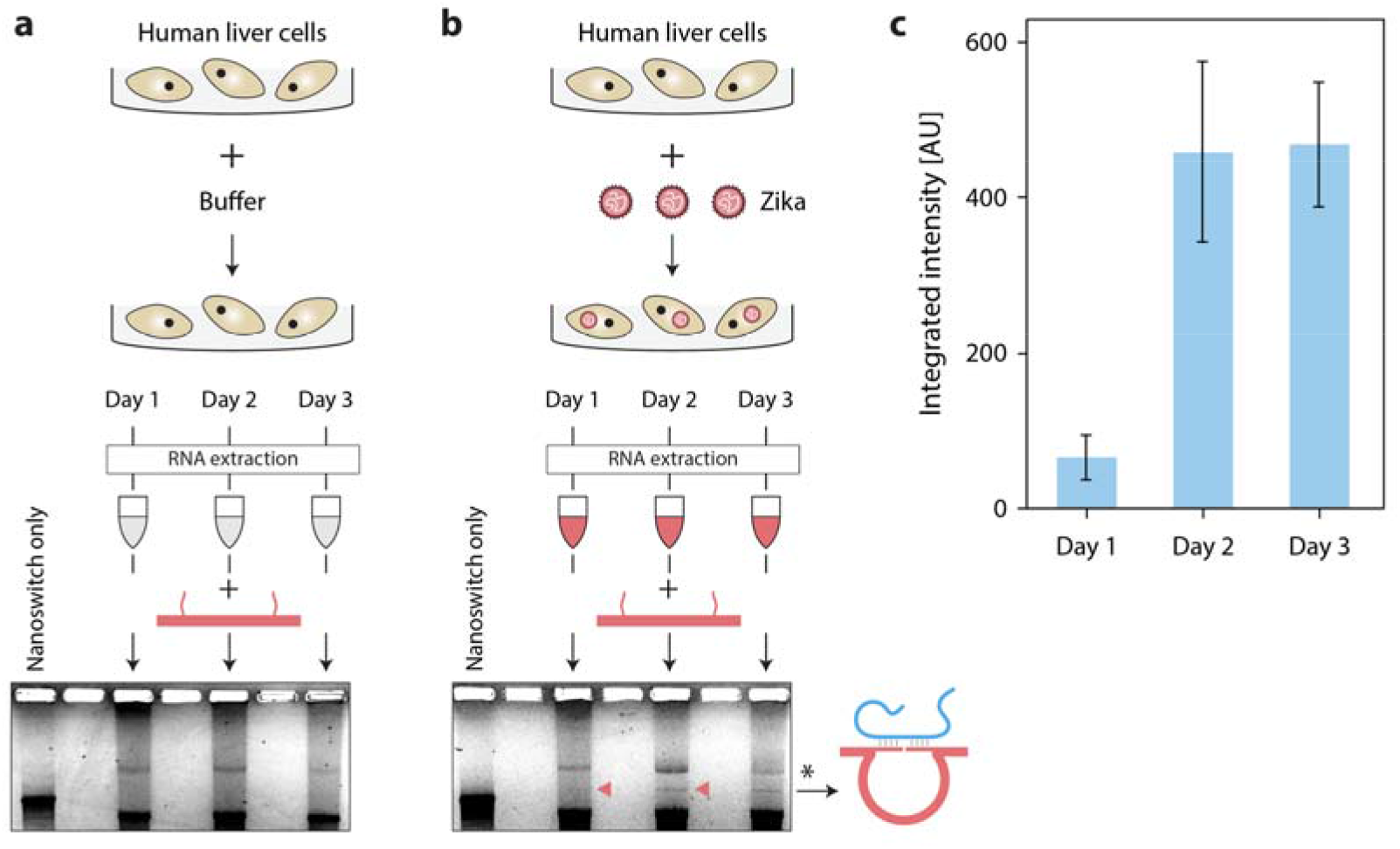
DNA nanoswitches directly detect ZIKV RNA from infected human liver cells. **(a)** RNA isolated from mock-infected Huh7 cells at 1, 2, and 3 days post infection show no ZIKV detection. (**b)** RNA isolated from Zika-infected Huh7 cells at 1, 2, and 3 days post infection shows increasing ZIKV detection over time, with red arrows denoting detection bands. * denotes a band of contaminating cellular DNA following RNA extraction. (**c)** Quantification of nanoswitch detection signal, with error bars representing standard deviation from triplicate experiments.

Moving toward clinical applications, we aimed to demonstrate detection of relevant levels of ZIKV RNA from biological fluids. ZIKV is present in the serum, urine, and other body fluids of infected patients.^39^ The viral loads can vary dramatically between individuals, body fluid, and post infection time,^6,7^ but are frequently in the subfemtomolar to femtomolar range, with ZIKV in human urine reported as high as 220 × 10^6^ copies/ml (365 fM).^33^ While our nanoswitch sensitivity for *in vitro* transcribed viral RNA in buffer approaches clinically relevant concentrations, detection from body fluids is further challenged by varying viral loads and by body fluids that can reduce the performance of the nanoswitches due to physiological conditions and nuclease activity.^26,40^ To overcome these potential difficulties, we investigated two independent solutions: 1) adding a pre-processing step to extract RNA from body fluids such as urine, or 2) adding an isothermal pre-amplification step. In the first approach, we spiked a clinically-relevant amount of *in vitro* transcribed ZIKV RNA into human urine and processed viral RNA extraction using a commercial RNA extraction kit. We then mixed the extracted RNA with the nanoswitches and demonstrated non-enzymatic, clinical level detection of the RNA at 0.28 pM (**Fig. 5a** and **Fig. S14**). In the second approach, we demonstrated that our detection can be coupled with other amplification approaches such as nucleic acid sequence-based amplification (NASBA).^41,42^ NASBA combines multiple enzymes and primers to achieve RNA amplification in a one-pot isothermal reaction (**Fig. S15A**). First, we showed feasibility of the amplification of ZIKV RNA by NASBA in water, followed by nanoswitch detection (**Fig. S15**). To mimic clinical samples, we spiked infectious ZIKV particles into either PBS or 10% human urine at clinical-levels (1.49 fM to 0.03 fM). From these samples our assay detected ZIKV RNA in ~5 hours (**Fig. 5b** and **Fig. S15**). We went one step further and showed that our assay can be performed using a commercially available bufferless gel cartridge (ThermoFisher E-gel) and imaged on a small and potentially portable gel reader (**Fig. S16**). With the help of NASBA amplification, the detection ability of our method has about 10,000-fold increase, from ~350 fM (**Fig. 5a**) to 0.03 fM (**Fig. 5b**) and the detection time was reduced from ~13 hours to ~5 hours.

**Fig. 5.**
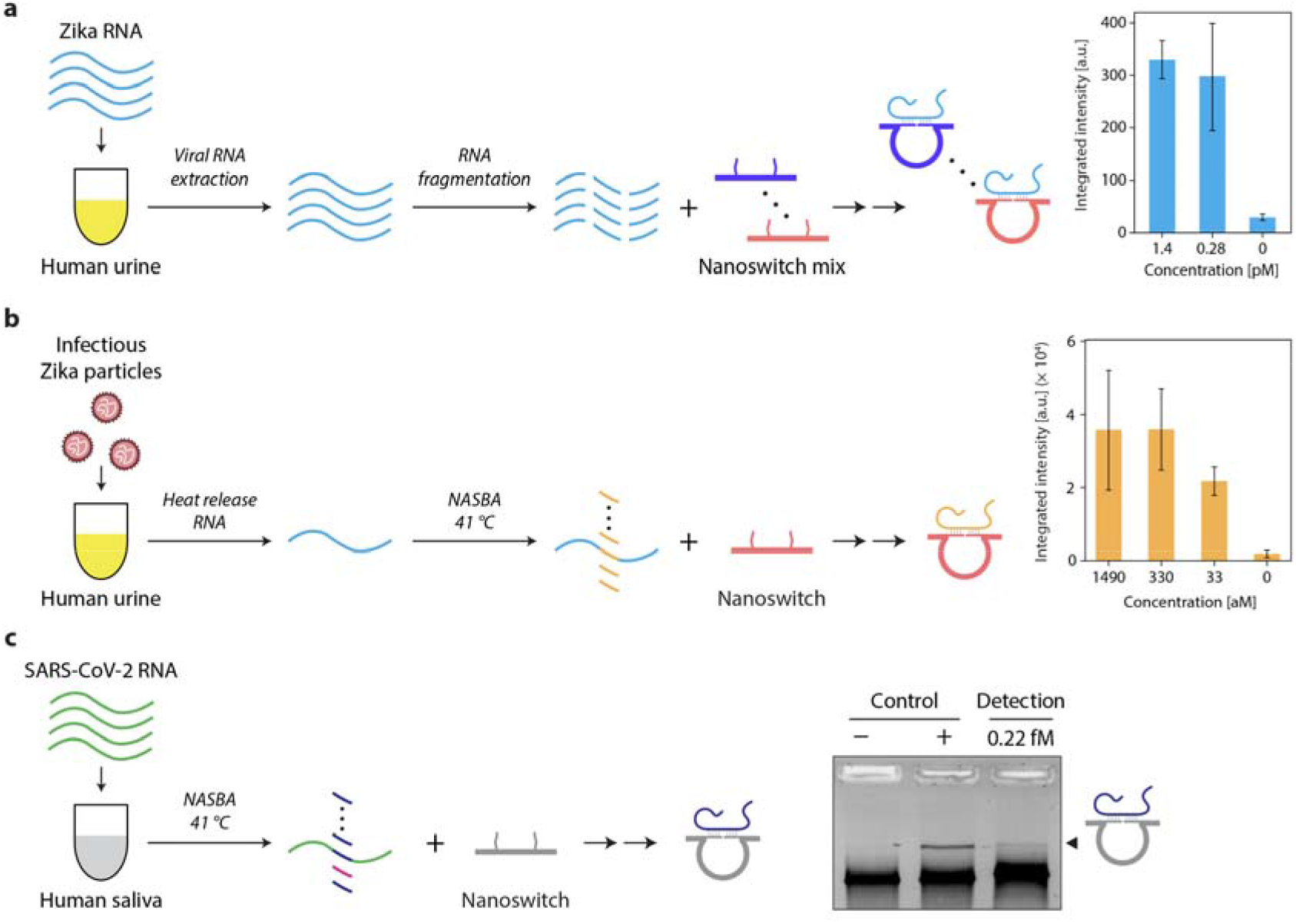
Prior extraction or pre-amplification of target RNA facilitates detection of ZIKV and SARS-CoV-2 RNA at clinically relevant levels in biofluids. **(a)** Positive identification of ZIKV RNA in spiked urine by first isolating *in vitro* transcribed target RNA using a commercially available viral RNA extraction kit, followed by direct, non-enzymatic detection using DNA nanoswitches. (**b)** Positive identification of ZIKV RNA from virus particles spiked into urine based on NASBA. **(c)** Positive detection of in vitro transcribed SARS-CoV-2 RNA in human saliva based on NASBA. Error bars represent standard deviation from triplicate experiments.

With the emerging outbreak of SARS-CoV-2 in January 2020, we took the opportunity to develop and test our DNA nanoswitches against the new virus. Following a similar strategy as for ZIKV, we identified a target region, developed nanoswitches, and used the NASBA strategy to detect a SARS-CoV-2 RNA fragment in 10% human saliva. We detected the fragment at a concentration of 0.22 fM (~100 copies/μl), which is around the clinical median (**Fig. 5c**).^43^ We compared this with our own RT-qPCR detection where we found a similar detection level (**Fig. S17**).

Taken together, we demonstrate that programmable DNA nanoswitches can be developed into a robust viral RNA detection platform, that is readily adaptable as we show in the detection of SARS-CoV-2. The platform has key advantages over existing methodologies in terms of selectivity and specificity, as shown in our experiments with ZIKV and closely related DENV, as well as two closely related ZIKV strains. Moreover, DNA nanoswitch viral RNA detection strategy has femtomolar detection limit without an RNA amplification step, and attomolar detection limit when used with amplification. These limits are within a clinically relevant range and therefore our DNA nanoswitch assay together with the bufferless gel cartridge presents a putative diagnostic assay for clinical detection of RNA viruses in low resource areas without significant laboratory infrastructure.

## Discussion

The functionality of our DNA nanoswitches is largely enabled by DNA nanotechnology, which has become a well-established field that uses DNA as a functional material to fabricate nanostructures.^44,45^ Biosensing is a particularly promising application of DNA nanotechnology,^46^ and reconfigurable DNA devices^47^ have been demonstrated for the detection of DNA,^47^ RNA,^48,49^ proteins,^50^ and pH.^51^ However, most designs are complex and require laborious readout with advanced microscopy that reduces their practicality. A few approaches have overcome this practicality hurdle to provide widely useful solutions to problems in biological imaging (e.g. DNA-PAINT in super-resolution microscopy^52^ and DNA scaffolds for NMR^53^ and cryo-EM^54^) and biosensing (e.g. detection of lysosomal disorders^55^ and mapping cellular endocytic pathways^56^). Our DNA nanoswitches take a reductionist approach, resulting in assays that are robust and sensitive, yet simple to adapt and do not require multiple steps or expensive equipment. With this work, we add viral detection to the existing suite of DNA nanoswitch assays that already includes protein^40^ and microRNA^27^ detection.

Our simple DNA nanoswitch-based assay for detection of viral RNA overcomes some limitations of currently available methods for clinical detection of RNA viruses in resource-limited areas. These challenges include 1) robust detection without enzymes or equipment, 2) maintaining low-cost and simplicity, and 3) providing specificity and versatility. Surprisingly, the current COVID-19 pandemic has shown us that these problems can affect rich countries as well, with most struggling to have testing outpace viral spread.

The intrinsically high signal of our nanoswitches is enhanced here with a new “target multiplication” strategy where we use viral RNA fragmentation to multiply the number of targets, and thus increase the signal intensity. Using this approach, we reached near-clinical levels of detection in urine without the use of enzyme-mediated amplification strategies. This is of significance because enzymes can be key drivers of assay cost and complexity due to requirements including cold storage/transportation, special buffers and reagents, and strict operating temperatures. These factors make enzymatic assays difficult for field use or for use in low resource areas without modern lab infrastructure. Despite these challenges, most currently available techniques rely on enzyme triggered amplification.^7,21,57^ For our assay, we demonstrated compatibility and dramatic signal improvement with an optional enzymatic preamplification step **(Fig. 5b-c)**. However, we believe that further improvements should enable complete coverage of the clinical range without enzymes. A 30 ml sample of urine from a ZIKV-infected patient would contain from 10^5^ to 10^9^ copies of viral RNA,^58^ theoretically surpassing our current detection limit. Efficient sample preparation using a viral RNA extraction kit **(Fig. 5a)**, for example, could facilitate use with our DNA nanoswitch assay.

Two key features of our approach are simplicity and low cost. Our DNA nanoswitches align with the goals of “frugal science” movement, where cost and accessibility to new technologies are valued alongside typical performance metrics.^59,60^ Our nanoswitches cost around 1 penny per reaction and can be stored dry at room temperature for at least a month, and could be delivered globally without transportation or biosafety concerns. The assay consists of few steps and can be performed in a matter of hours with limited laboratory needs (**Fig. S18**). Our assay uses a readout by gel electrophoresis, which is relatively inexpensive and already part of the workflow in many labs, which is comparatively simpler than many nanotechnology-based assays involving multiple incubation and wash steps. Improvements to the signal readout could potentially help make this approach even more lab independent. Successful detection with a commercially available bufferless gel system **(Fig. S16)** takes us a step closer to enabling field deployment of our assay, and sample preparation could be aided by other frugal science approaches such as the “paperfuge”^61^ and low-cost thermal cycler.^62^ If purified viral RNA is used in NASBA, the entire detection could be shortened to two hours with a 30 min NASBA step^21^ and a 1 hour nanoswitch detection assay.^27^

The programmability of our system makes it versatile for a wide variety of viruses including ZIKV, DENV, and SARS-CoV-2 as we have shown. These can be detected with high specificity as we showed for ZIKV and DENV **(Fig. 3a-b)** and for different strains of ZIKV **(Fig. 3c-d)**, even in a multiplexed fashion. Here we have focused on single-stranded RNA viruses but assays for other RNA or DNA viruses could likely be developed similarly. The fast construction and purification processes can facilitate rapid production of DNA nanoswitches to detect an emerging viral threat, potentially in as little as 1-2 days from knowledge of the target sequences, limited mostly by oligo synthesis turnaround time (**Fig. 1b**). Due to the low cost of the test, our assay could also be useful for monitoring viral progression over time in patients, or for testing potentially infected insects or animals. Therefore, with future optimization towards point-of-care clinical applications in resource-limiting environments, the platform we describe here has a potential to improve accuracy and ease of diagnosis in humans, non-human vectors, and other animals. Ultimately this can enhance our ability to control spread of infection and more rapidly respond to emerging viral threats including the COVID-19 pandemic, and work toward a reduced death toll and economic burden.

## Methods

### Construction and purification of nanoswitches

Oligonucleotides were purchased from Integrated DNA Technologies (IDT) with standard desalting, and the full sequences of all strands is listed in Supporting information **Table S1 to S10**. Nanoswitches were constructed as described previously.^24,27^ A genomic single-stranded DNA (New England Biolabs M13mp18) was linearized using targeted cleavage with BtsCI restriction enzyme. The linearized ssDNA was then mixed with a molar excess of an oligonucleotide mixture containing backbone oligos and detectors, and annealed from 90°C to 20°C at 1 °C min^-1^ in a T100™ Thermal Cycler (Bio-Rad, USA). Following construction, the nanoswitches were purified using liquid chromatography (LC) purification^29^ to remove excess oligonucleotides. The concentration of purified nanoswitches were determined by measuring A260 absorbance with a Thermo Scientific NanoDrop 2000.

### In vitro transcription (IVT) of viral RNA

Plasmids containing the full-length ZIKV (Cambodia FSS13025 strain; pFLZIKV) and DENV-2 (strain 16681, pD2/IC-30P) cDNAs were gifts from Dr. Pei-Yong Shi (University of Texas Medical Branch) and Dr. Claire Huang (Centers for Disease Control), respectively.^30,63^ pFLZIKV was linearized with ClaI (New England Biolabs, NEB), and pD2/IC-30P was linearized with XbaI (NEB). Digested plasmids were extracted with phenol:chloroform:isoamyl alcohol and then precipitated. Linearized plasmids were *in vitro* transcribed (Thermo Fisher Scientific) and the resulting viral RNA cleaned by MEGAclear Transcription Clean-Up Kit (Thermo Fisher Scientific). We followed the protocols of these two kits except that we did not heat the purification column in the elution step of the viral RNA because we noticed that high temperature can result in degradation of the viral RNA.

### Viral RNA fragmentation test

Viral RNA was fragmented by using 10×Fragmentation buffer (NEB) and the recommended protocol. Briefly, the ZIKV RNA obtained from *in vitro* transcription was mixed with fragmentation buffer (1× final) and then incubated at 94°C in a thermal cycler for 1, 3, 6, or 9 min. Then RNA fragmentation analyzer (Agilent, model 5003) was used to quantify the length distribution of RNA fragments by using the DNF-471 Standard Sensitivity RNA Analysis Kit (**Fig. 2b**).

### Cell culture, ZIKV infections and total RNA extraction

Human hepatocarcinoma (Huh7) cells were maintained in Dulbecco’s modified Eagle’s medium (DMEM) (Life Technologies) supplemented with 10% fetal bovine serum (FBS; VWR Life Science Seradigm), 10 mM non-essential amino acids (NEAA; Life Technologies), and 5 mM L-glutamine (Life Technologies). Cells were passaged once every three days and maintained at 37°C with 5% CO2. Twenty-four hours prior to infection Huh7 cells were seeded into tissue culture plates. The following day, one plate was counted. The other two plates were used for mock- and ZIKV-infection, where cells were infected at a multiplicity of infection of one. The original Cambodia and Uganda (MR766) ZIKV stocks were a generous gift from Dr. Brett Lindenbach (Yale School of Medicine). RNA from mock- and ZIKV-infected cells was harvested by removing media from the cells was removed and then washing the cell monolayer washed once with ice-cold PBS. Hereafter, each tissue culture plate was lysed in 1 mL TRIzol (Invitrogen) and total RNA extracted per the manufacturer’s instructions. For experiments using ZIKV infectious particles in PBS/urine, cell culture media from mock- and ZIKV-infected cells was collected and stored at −80 °C. The number of infectious particles was determined by plaques assay, as described previously.^38^

### DNA nanoswitch detection

The total detection sample volume was 10 μl with 10 mM MgCl2, 1×PBS, 1.5-2 μl nanoswitch at 0.1 nM. The detection incubation was finished in a thermal cycler with a thermal annealing from 40 °C to 25 °C at −1 °C min^-1^ or room temperature (e.g. the NASBA related detections). Before loading to the gel, the sample was stained by GelRed (Biotium Inc.) at 1× concentration (3.3×GelRed for total RNA related detection) and mixed with 2 μl 6× Blue loading dye.

### Viral RNA detection sensitivity test

For the experiments in **Fig. 2f** and **Fig. S8**, first, all nanoswitches were purified by LC and then their concentrations were determined by measuring A260 absorbance with a Thermo Scientific NanoDrop 2000. Nanoswitch mixtures were made by mixing nanoswitches in equimolar concentrations. The detection reaction volume is 10 μl with nanoswitch (0.1 nM), MgCl_2_ (10mM), 1×PBS and blocking oligos (200 nM). The blocking oligos are short oligos (14 nucleotides) that can prevent the binding of target RNA to the inner surface of plastic tubes.^27^ The detection incubation was conducted in a thermal cycler with a thermal annealing from 40 °C to 25 °C for about 12 hours.

### Detection of viral RNA from total RNA

First, 500 ng total RNA extracted from uninfected/infected cells was fragmented at 94°C for 9 minutes in 1x fragmentation buffer. Fragmented total RNA was then mixed with nanoswitches (1 nM), MgCl_2_ (10 mM) and PBS (1×), and the mixture was made up to 10 μl with nuclease-free water. Samples were then incubated in a thermal annealing ramp from 40 °C to 25 °C at 1 °C min^-1^. After the incubation, samples were stained with GelRed (Biotium Inc.) at 3.3× concentration and incubated at room temperature for 30 min. Before loading the gel, 2 μl of 6x blue loading dye was mixed with each sample, and 10 μl sample was loaded to each well. Samples were run in a 0.8% agarose gel at 65-75 V for about 70-90 minutes in the cold room.

### Detection of viral RNA extracted from urine

For the detection of viral RNA extracted from urine, we first spiked the ZIKV RNA with blocking oligos (200 nM) into 200 μl human urine to mimic the clinic sample. Then, we used Quick-RNA Viral Kit (Zymo research) to extract the viral RNA from it. Different amount of ZIKV RNA were tested (**Fig. 5a**). Here, the amount of human urine can be scaled up as needed according to the protocol of the kit. Finally, the viral RNA was eluted from the filter column by using 15 μl nuclease-free water. Then 5 μl extracted RNA was fragmented at 94°C for 9 minutes by using 0.2× fragmentation buffer (NEB) before conducting the nanoswitch detection. Here, we lowered the use of fragmentation buffer in the consideration of the small amount of RNA in the extracted sample as we noticed that too much fragmentation buffer could destroy the DNA nanoswitches (data not shown).

### Isothermal amplification by NASBA

First, we employed the classic NASBA protocol^41^ to prove the concept (**Fig. S15**). The 25 μl one-pot reaction contained 3 μl RNA sample at various concentrations, 0.4 μM forward and reverse primers, 8 U AMV Reverse Transcriptase, 50 U T7 RNA Polymerase, 0.1 U RNase H, 40 U RNase Inhibitor (NEB, Murine), 2 mM NTP mix, 1 mM rNTP mix, 12 mM MgCl2, 40 mM Tris-HCl, 42 mM KCl, 5 mM Dithiotreitol (DTT), 15%(v/v) dimethyl sulfoxide. The primers were chosen from reference ^21^. The sample was incubated at 41 °C for 2 hours in the thermal cycler and followed by heating at 94 °C for 10 minutes to deactivate all enzymes. Three microliters of the sample were used in the following DNA nanoswitch detection assay in PCR tubes with 10 μl final volumes. After mixing with the DNA nanoswitch and reaction buffer, the mix was incubated at room temperature for two hours. GelRed (Biotium Inc.) at 1×concentration was added to the detection samples before loading to the 0.8% agarose gel. The gel was run at room temperature for 45 minutes at 75 volts.

For the ZIKV related NASBA experiments, first we spiked the ZIKV infectious particles (started at 1.96 fM of particles) into PBS or human urine. Then samples were diluted into different concentrations 1.49, 0.33, and 0.03 fM with 1xPBS or 10% human urine. Subsequently, blocking oligo (200 nM) was added and the viral RNA was released by heating the samples at 94 °C for 3 minutes. For the human urine samples, RNase Inhibitor (NEB, Murine) was also added at concentration of 2 U/μl before heating. After cooling down at room temperature, 0.5 μl RNase Inhibitor at 40 U/ul (NEB, Murine) was added to 10 μl human urine sample to protect the viral RNA. Total volume of each NASBA reaction was scaled down to 6 μl which contains 1.25 μl Enzyme COCKTAIL (NEC 1-24), 2 μl 3× buffer (NECB-24), 0.48 μl NTPs mix at 25 mM, 0.3 μl dNTPs mix at 20 mM, 0.2 μl two primers mix at 10 uM, 0.2 μl RNase Inhibitor with 40 U/ul (NEB, Murine) and 1.57 μl sample with viral RNA. The sample was incubated at 41 °C for 2 hours in the thermal cycler and followed by heating at 94 °C for 10 minutes to deactivate all enzymes. Then 1 μl of the NASBA sample was used in the following DNA nanoswitch detection assay in PCR tubes with 10 μl final volumes. The assay was finished by incubating at room temperature for two hours. After mixing with GelRed (Biotium Inc.) at 1×concentration and 2 μl 6× blue loading dye, the detection samples were loaded to the 25 ml 0.8% agarose gel which was run in 0.5×Tris-Borate-EDTA (TBE) buffer at 75 V at room temperature for 45 minutes.

### Detection of SARS-CoV-2 RNA

A gBlock gene fragment of the SARS-CoV-2 RNA segment ^64^ was purchased from IDT (**Table S9**). Then, PCR amplification (Qiagen, Taq PCR Core Kit) was used to create more copies with a T7 promoter that was added to the 5’ end of the forward primer. Afterword, dsDNA template was cleaned by the QIAquick PCR Purification Kit (Qiagen). Finally, the SARS-CoV-2 RNA was obtained by in vitro transcription (NEB, HiScribe™ T7 High Yield RNA Synthesis Kit) and cleaned by MEGAclear Transcription Clean-Up Kit (Thermo Fisher Scientific). The RT-PCR detection was conducted by using the Luna Universal One-Step RT-qPCR kit from NEB and following its protocol. 1.5 ul RNA sample was used in 20 ul reaction mix.

### Gel imaging and analysis

The detection samples were run in 0.8% agarose gels unless otherwise noted, cast from molecular biology grade agarose (Fisher BioReagents) dissolved in 0.5× TBE buffer. Typical running conditions were 75 V for 45 to 70 minutes at room temperature or cold room. Samples were mixed with a Ficoll based blue loading dye prior to loading. Imaging was completed on a Bio-Rad Gel Doc XR+ imager with different exposure time based on the brightness of the detection bands. The detection efficiency was analyzed using included Image Lab software (**Fig. 2e**). The profiles of detection bands were obtained in ImageJ ^65^ and then their integrated intensity were obtained by using the peak analysis function in Origin (OriginLab Corporation), such as the data presented in **Fig. 2f, 4c and 5a-b**. Detailed analysis procedure can be found in our previous publications^27^. For the E-gel related experiments, we used Invitrogen E-gel agarose system (Thermo Fisher Scientific) and its precast agarose gel (1.0%, SYBR stained). 10 μl of nanoswitch detection sample was loaded to each lane and the gel was run at 48 volts for 1 hour at room temperature. Since the E-gel system does not allow user control of the voltage, we used an external power supplier connected with the negative and positive electrodes of the precast agarose gel to supply 48 volts.

## Acknowledgements

We thank Andy Berglund, Carl Shotwell, and Jared Richardson for facilitating use of the RNA fragment analyzer; Darren Yang and Wesley P. Wong for assistance and suggestion on adapting the E-gel system with the nanoswitches; Gabriele Fuchs and Bijan K. Dey for providing comments and suggestions on the experiments and manuscript.

## Author contributions

L.Z., K.H., and C.T.P. designed the experiments. L.Z., A.R.C., J.A.P., S.C., O.L., C.C., and K.H. performed the nanoswitch experiments. G.B., P.B., and C.T.P. carried out the cell culture and total RNA extraction from infected cells. L.Z. developed the tool in Matlab and wrote the first draft of the manuscript. L.Z., A.R.C., J.A.P., C.T.P., and K.H. co-wrote later drafts of the manuscript.

## Data and materials availability

All data needed to evaluate the conclusions in the paper are present in the paper and/or in the Supplementary Materials. Additional data related to this paper may be requested from the authors.

