## Supplemental Information for "Programmable low-cost DNA-based platform for viral RNA detection"

**Contents**

***Figures***

**Fig. S1.** In vitro transcription (IVT) of viral RNA.

**Fig. S2.** Fragmentation analysis ZIKV RNA.

**Fig. S3.** Optimization of the detection arm length.

**Note S1.** Choosing the detection targets of viral RNA.

**Fig. S4.** Considerations for choosing target sequences of viral RNA.

**Fig. S5.** Schematic showing assembly of DNA nanoswitch.

**Fig. S6.** Graphical user interface (GUI) for obtaining potential viral RNA targets.

**Fig. S7.** Analysis of the 18 DNA nanoswitches designed for ZIKV RNA detection.

**Fig. S8.** An example of gel image of the 18 mixed nanoswitches detection sensitivity test.

**Fig. S9.** Detection sensitivity test of single nanoswitch.

**Fig. S10.** Analysis of the 12 DNA nanoswitches designed for DENV RNA detection.

**Fig. S11.** Tuning the loop size of DNA nanoswitch.

**Fig. S12.** Targets and a gel image of specificity test with Cambodia and Uganda strains of ZIKV.

**Fig. S13.** Detection of ZIKV RNA in total RNA extracted from human liver cells.

**Fig. S14.** Gel images of the ZIKV RNA detection in samples mimicking the urine of patients.

**Fig. S15.** Detection of ZIKV RNA based on pre-amplification with NASBA.

**Fig. S16.** Portable e-gel system for detection of ZIKV RNA based on pre-amplification with NASBA.

**Fig. S17.** Detection of SARS-CoV-2 RNA in human saliva.

**Fig. S18.** Summary of the viral RNA detection procedure by using DNA nanoswitches.

***Tables***

**Table S1.** A ZIKV RNA target sequence from a literature^5^ and its corresponding detection arm ssDNA (experiments in Fig. 2c).

**Table S2.** Target sequence and different lengths of detection arm ssDNA (15, 14, 13, 12, 11, 10nt) for optimizing the design of nanoswitch (experiments in Fig. S3).

**Table S3.** The eighteen target sequences and corresponding detection arm ssDNA oligos for the detection of ZIKV RNA (experiments in Fig. 2e, 2f, 3a, 3b, 3d, 5a and S7, S8, S9).

**Table S4.** The twelve target sequences and corresponding detection arm ssDNA oligos for the detection of DENV RNA (experiments in Fig. 3a, S10).

**Table S5.** Variable oligos for constructing nanoswitches with different loop sizes (experiments in Fig. 3b, 3d).

**Table S6.** Target sequences and the corresponding detection arm ssDNA for the ZIKV and DENV multiplexing test (experiments in Fig. 3b).

**Table S7.** Target sequences and the corresponding detection arm ssDNAfor the ZIKV Cambodia and Uganda specificity test (experiments in Fig. 3d).

**Table S8.** Amplified region of ZIKV RNA, primers, targets and corresponding detection arm ssDNA used in NASBA (related experiments in Fig. 5b, S15, S16).

**Table S9.** DNA template, primers, target and the corresponding detector ssDNA for SARS-CoV-2 RNA detection.

**Table S10.** Backbone and basic variable oligos for the construction of nanoswitches and other oligos.

***Other supporting information***

Matlab code for selecting viral RNA targets for DNA nanoswitch assay (**File 1**)

Instruction for the Viral RNA detection target tool (PDF)

***References***

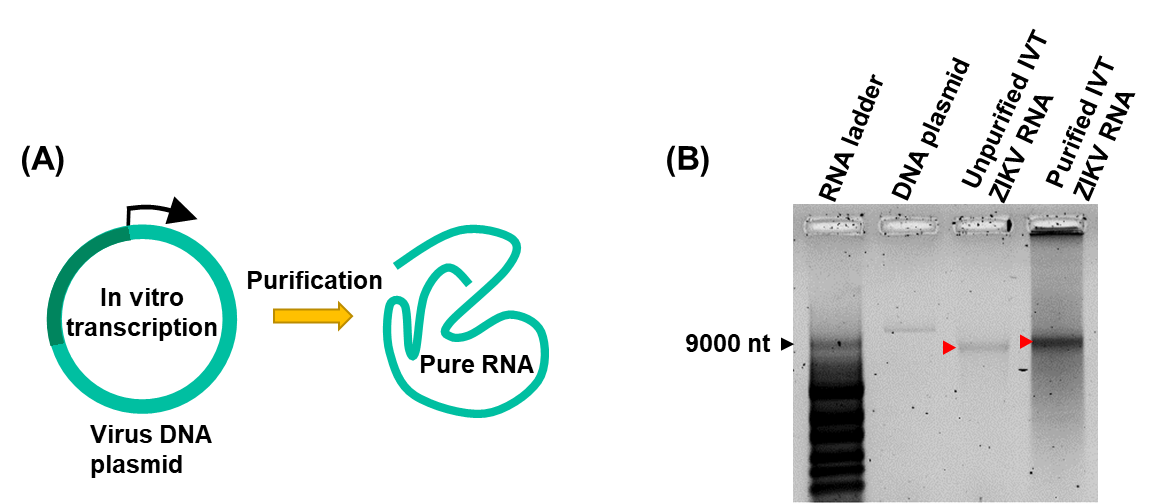

### **Fig. S1. In vitro transcription (IVT) of viral RNA.** (**A**) Schematic of in vitro transcription reaction. Plasmids containing the full-length infectious cDNA clone of either the ZIKV or DENV genomes were linearized, *in vitro* transcribed, followed by purification of the RNA product. (**B**) Integrity of *in vitro* transcribed ZIKV RNA was analyzed by electrophoresis in a native 0.8% agarose/TBE gel. Red arrow indicated the band of ZIKV RNA. Note: IVT and purification were finished by using the MEGAscript™ T7 Transcription Kit and MEGAclear™ Transcription Clean-Up Kit from Thermo Fisher Scientific. We followed the protocols of these two kits except that we didn’t heat the purification column in the elution step of the viral RNA because we noticed that high temperature could result in degradation of the viral RNA.

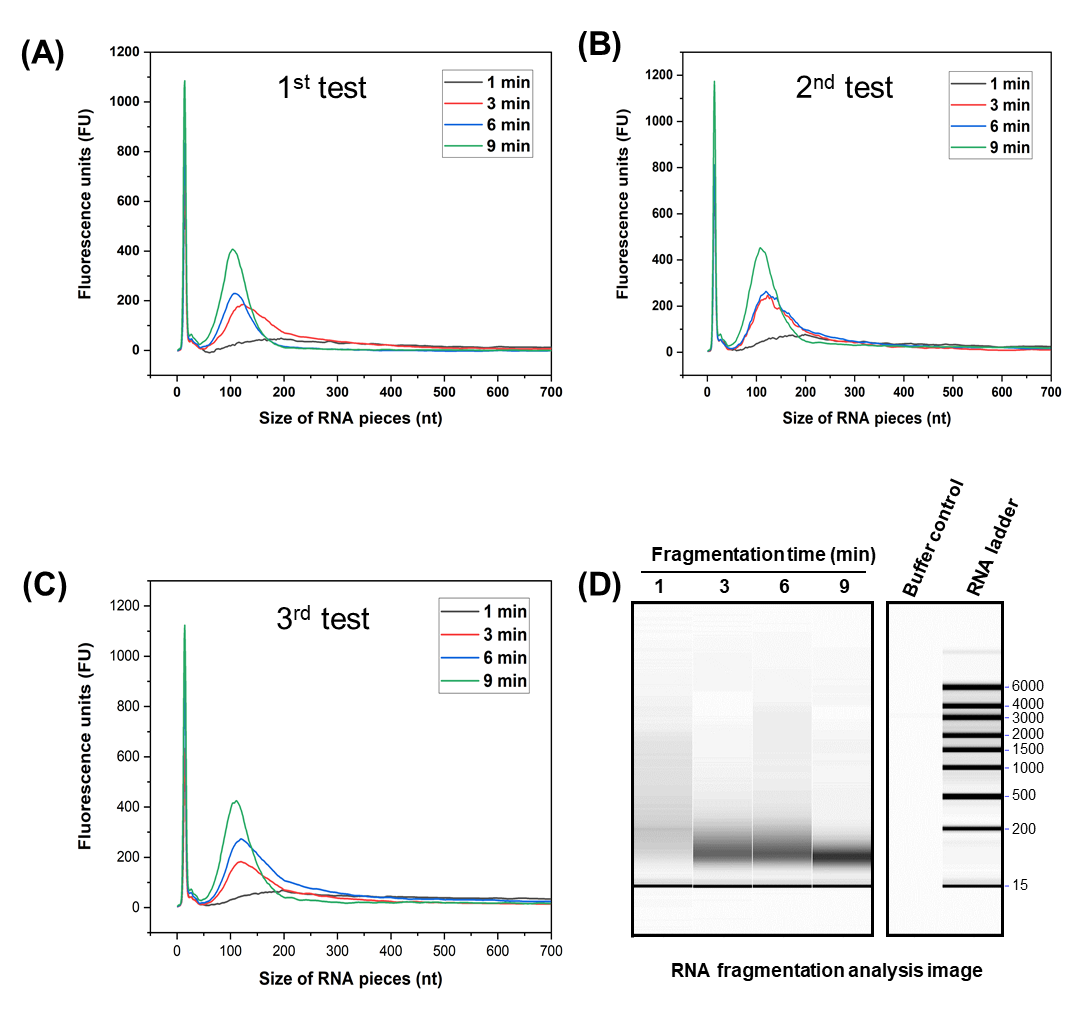

### **Fig. S2. Fragmentation analysis ZIKV RNA.** (**A**, **B**, **C**) Triplicate results of the ZIKV RNA fragmentation. *In vitro* transcribed ZIKV RNA was fragmented at 94 °C using the RNA fragmentation buffer from New England Biolabs for 1, 3 6 and 9 minutes. (**D**) An example of fragmentation gel image from the RNA fragmentation analyzer machine showing optimal fragmentation abundance and size following 9 minutes of fragmentation.

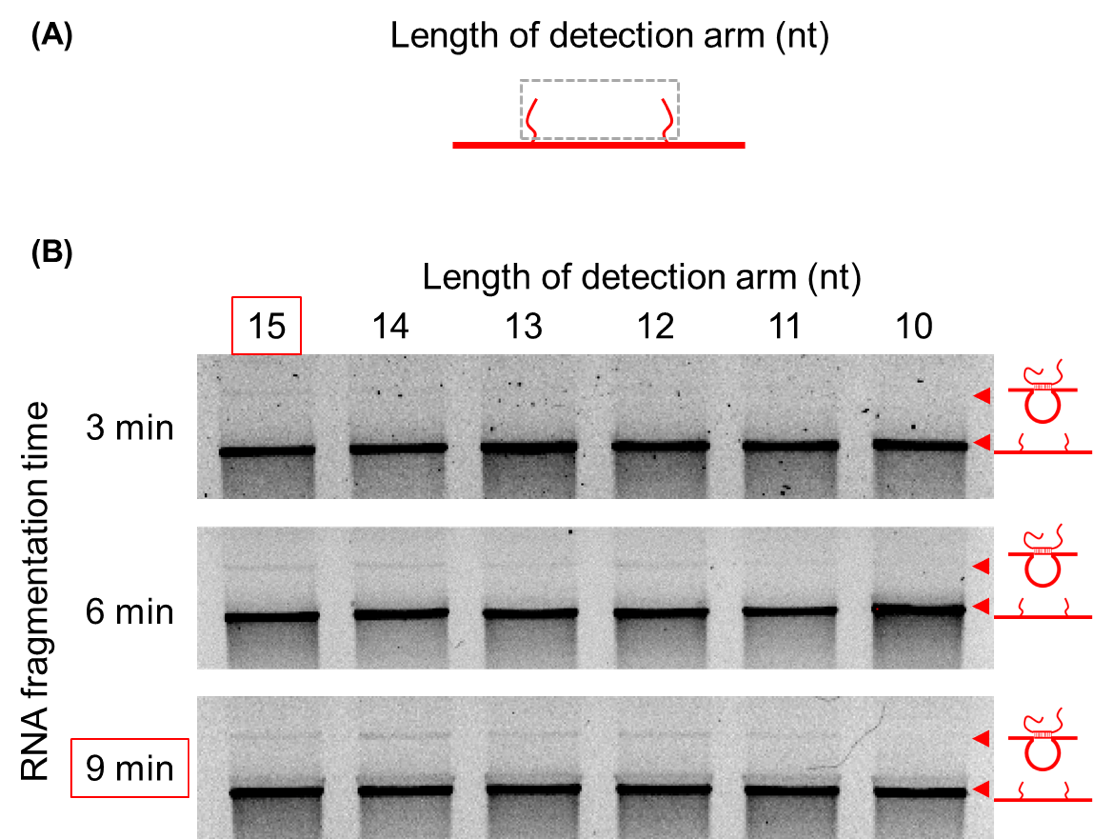

### **Fig. S3. Optimization of the detection arm length.** (**A**) Schematic of the DNA nanoswitch. (**B**) Nanoswitches with detector oligonucleotides of different lengths (10-15 nucleotides long) were incubated with *in vitro* transcribed ZIKV RNA that was fragmented at 94 °C with the NEB fragmentation buffer for 3, 6 and 9 minutes. An example 0.8% agarose/TBE gel image showing detection of ZIKV RNA. These results revealed optimal detection of ZIKV RNA following 9 minutes of RNA fragmentation and with a nanoswitch containing a 15-nucleotide detector arm length. The nanoswitch used in this experiment is the third nanoswitch in **Table S3**.

### **Note S1 Choosing the detection targets of viral RNA**

We first determined the target length to be 30 nt based on the detection test results of **Fig. S3**. The ZIKV genome is approximately eleven thousand nucleotides. Within the genome, the RNA can form very stable secondary structures that could inhibit detection by the DNA nanoswitches. We excluded those regions based on the minimum free energy (MFE), a parameter used to indicate the stability of the secondary structures of potential targets. The lower the MFE, the more stable the secondary structure will be. In addition, it is helpful to have the target sequence with a relatively high GC-content that can enhance the hybridization between the ssDNA detection arms of the nanoswitch and the target RNA. To ensure specificity of our target, we also examined sequence similarity between ZIKV and DENV genome sequences and eliminated those sequences with high alignment scores. When investigating different strains of the same virus the sequence similarity is high and it is effective to pick target regions with as many different nucleotides within the region of interest. The detailed procedure for choosing target sequences is described below and the corresponding tool developed in Matlab can be found in **File S1**.

**Step 1:** Create the target pool based on the detection region length, GC-content (≥35%) and minimum free energy (≥-2 kcal/mol). The minimum free energy was calculated by using the Matlab function: rnafold(seq).

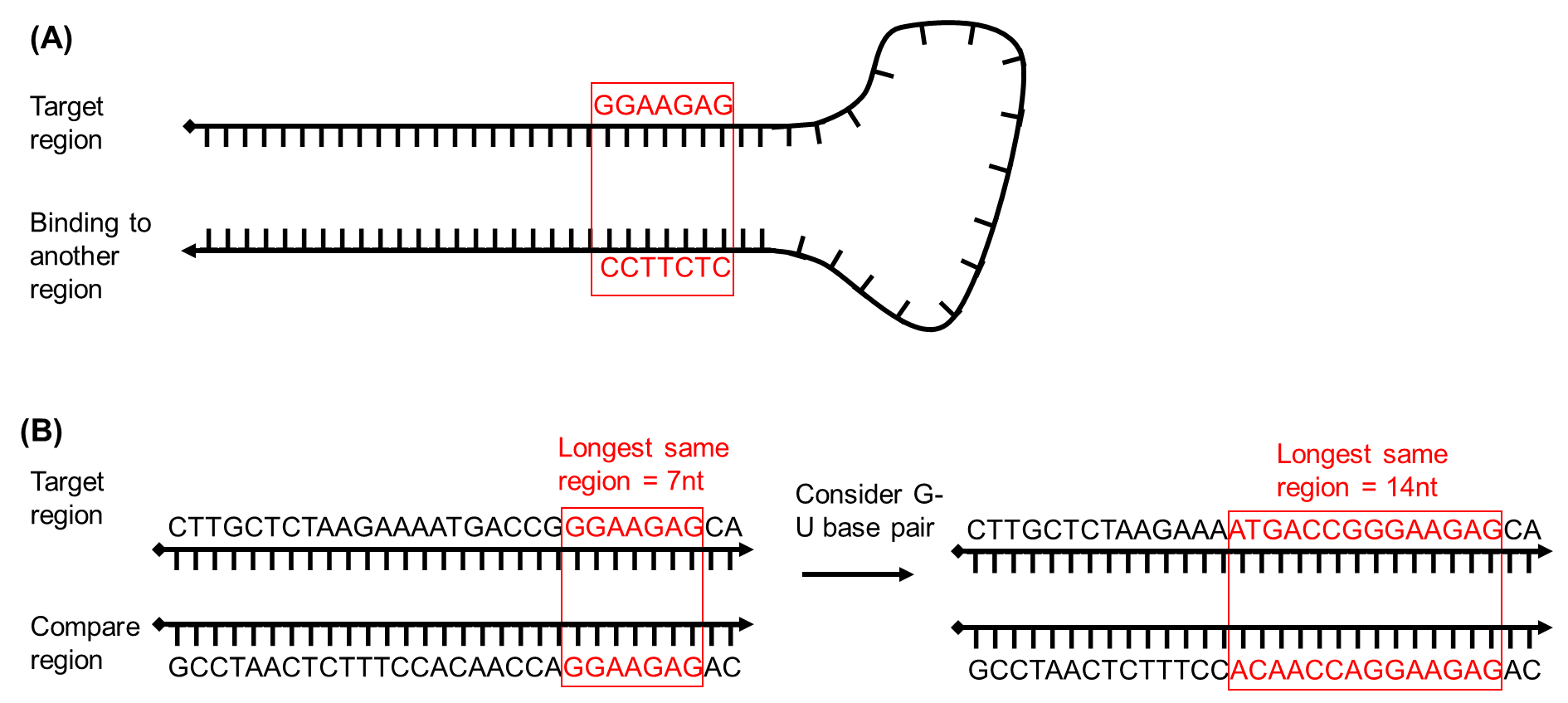

### **Fig. S4. Considerations for choosing target sequences of viral RNA.** (**A**) Schematic of self-binding and formation of a stable secondary structure that should be excluded as a target sequence. (**B**) An example of two comparable targets when G-U base pairing is taken into consideration.

**Step 2:** Check the potential for strong self-binding within the viral RNA sequence (Fig. S4A). The similarity between two target regions was quantified from the alignment score obtained by the Matlab function: nwalign(Seq1,Seq2). Higher alignment score corresponds to higher similarity. When comparing two regions, the program also computed the number of same nucleotides and the length of longest adjacent same nucleotides (Fig. S4B). Because G-U base pairing plays an important role in the formation and stabilization of RNA secondary structures, here we also took G-U base pair into account (Fig. S4B). Then, we eliminated the pair of targets that have the length of same adjacent nucleotides longer than 13 nt when G-U base pair is considered.

### **Step 3:** Check the similarity of targets obtained in Step 2 with the DENV RNA sequence (Dengue virus serotype 2, strain Thailand 16681; Genbank accession NC001474)^1^ and remove the targets that could result in cross detection with DENV. Here the criteria were that the length of longest adjacent same nucleotides should be no longer than 15nt and 20nt when G-U base pairing is considered.

**Step 4:** Check the similarity of targets with the complementary sequence of M13 (p7249), which is used to construct the nanoswitch. This avoids the binding of the ssDNA detection arms with the backbone ssDNA (Fig. S5). Here the criteria are that the length of longest adjacent same nucleotides on both half side (13 nt) of the target should be no longer than 6nt based on our previous research^2^.

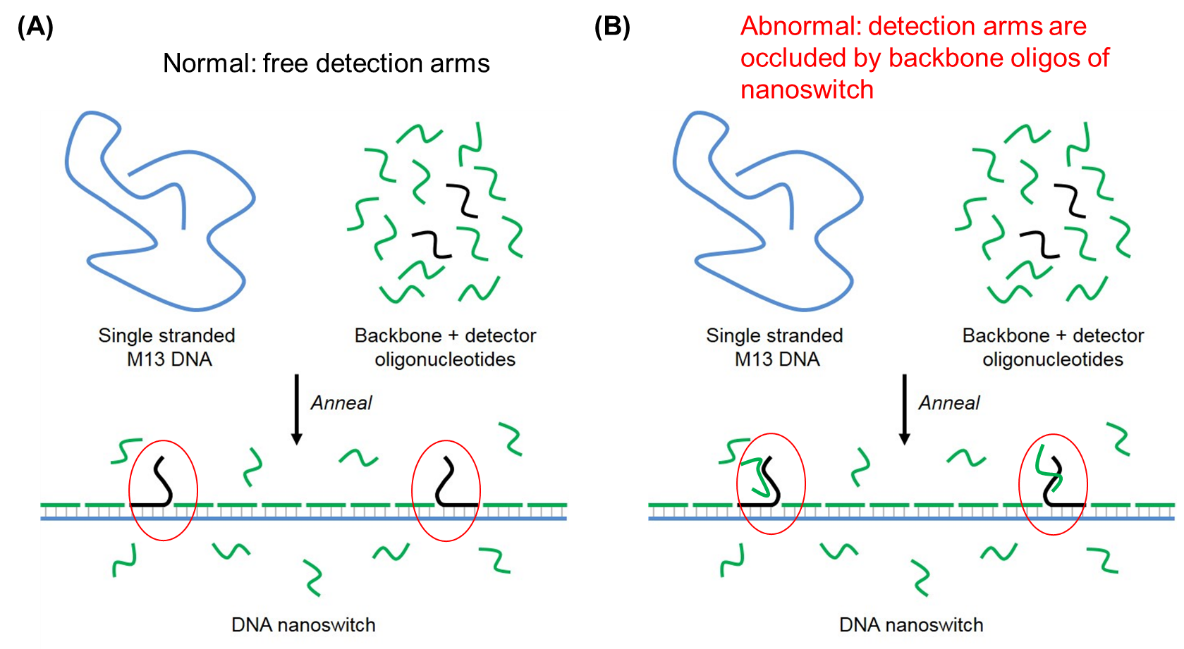

### **Fig. S5. Schematic showing assembly of DNA nanoswitch.** (**A**) Single stranded M13 DNA is annealed with backbone oligonucleotides and detectors specific to the RNA target. Normal assembly results in the detection oligonucleotides have free detection arms. (**B**) In contrast, abnormal assembly of the DNA nanoswitch may result when excess backbone oligonucleotides interact with the detector oligonucleotides and occlude the detection arms thus blocking recognition of target RNA.

**Step 5:** Pick targets from the final list to ensure the distance between them is longer than 50 nt. The performance of the nanoswitches could be first verified by positive control experiment that uses corresponding ssDNA as the target. The Matlab code with GUI is also provided in the supporting material with instructions for users (see **Fig. S6.** and **Instruction for the Viral RNA detection target tool.pdf**).

**Step 6 (Optional):** If different strains of the same virus are required to be detected, then the targets should satisfy the requirement that there should be more than 5 mutations between the targeted regions from the two strains. In addition, the position of mutation nucleotides should be near the middle of the detection arm. As the mutation number and position have higher priority that the factors discussed in steps 2-5, the potential targets are screened and picked from the target pool obtained in step 1.

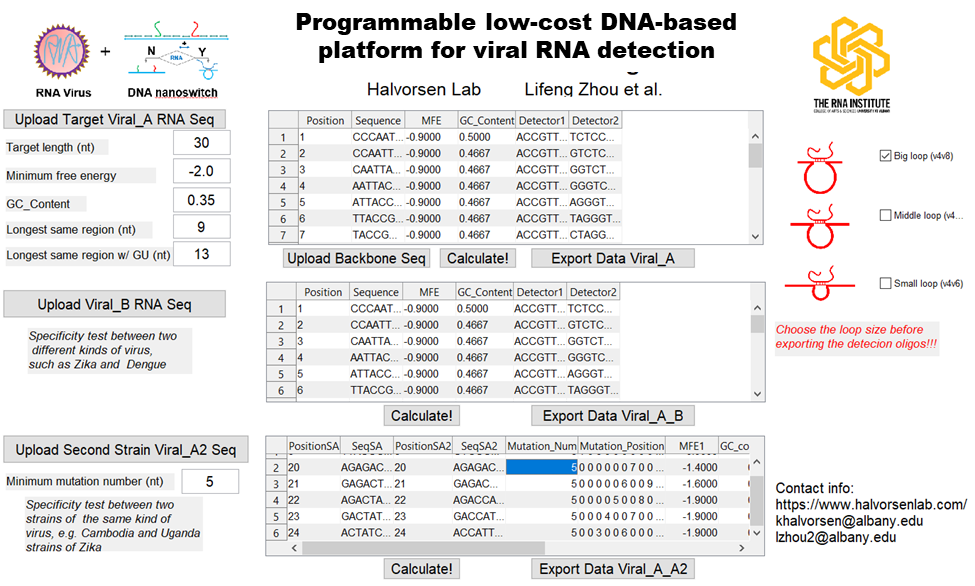

**Fig. S6. Graphical user interface (GUI) of the tool for obtaining potential targets of viral RNA for DNA nanoswitch detection.** More requirements could be added to the procedure and the Matlab code can be easily customized to obtain the desired target regions of viral RNAs (See **Instruction for the Viral RNA detection target tool.pdf** and **File S1**).

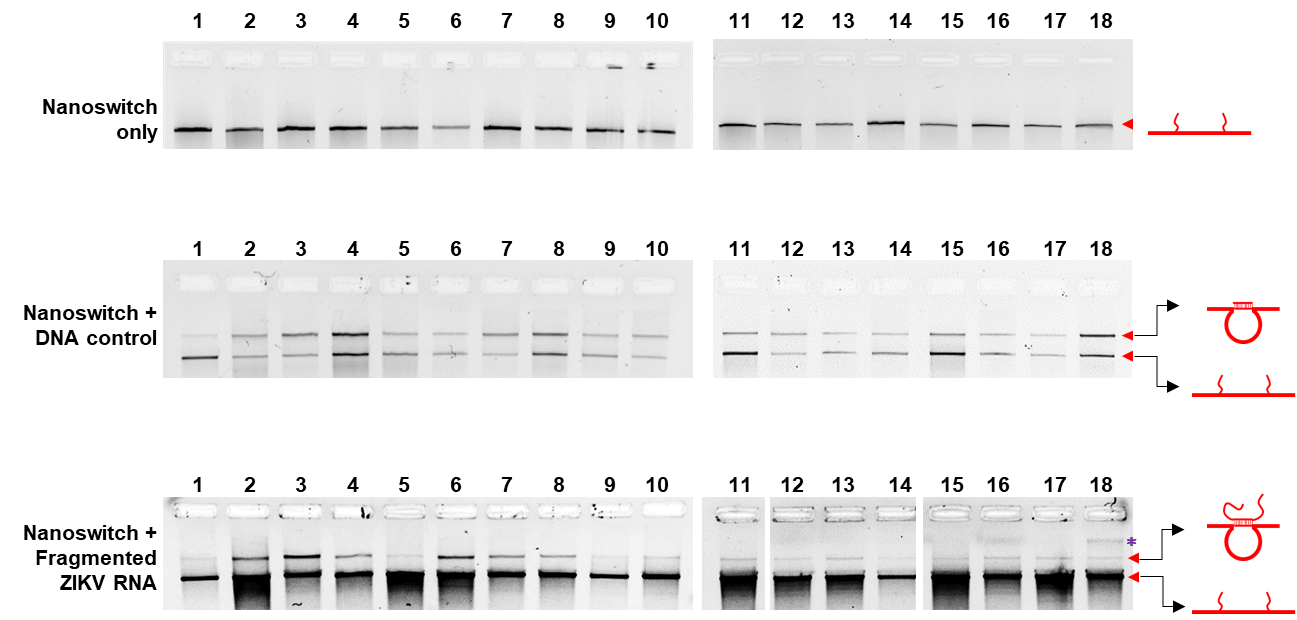

### **Fig. S7. Analysis of the 18 DNA nanoswitches designed for ZIKV RNA detection.** Top panel shows the negative control test of just the different DNA nanoswitches. The middle panel shows the positive control of complementary ssDNA (2 nM) annealed with the corresponding nanoswitch. The bottom panel shows detection of ZIKV RNA by individual DNA nanoswitches. We used 5 ng of fragmented *in vitro* transcribed ZIKV RNA to test the nanoswitches.

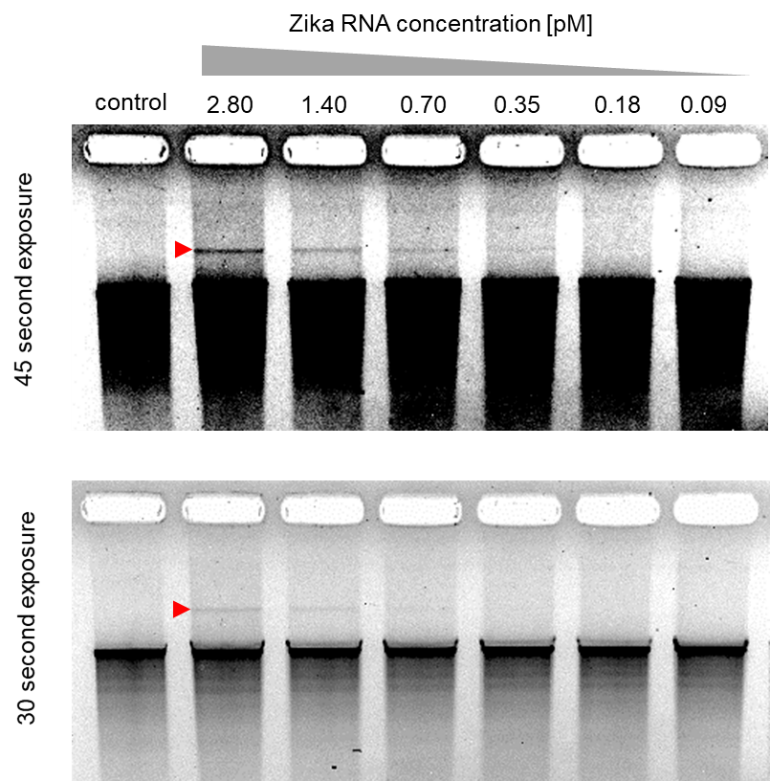

### **Fig. S8. An example of gel image of the 18 mixed nanoswitches detection sensitivity test**. This is the representative gel shown in **Fig. 2f** (45 second exposure is shown at the top and 30 second exposure at the bottom).

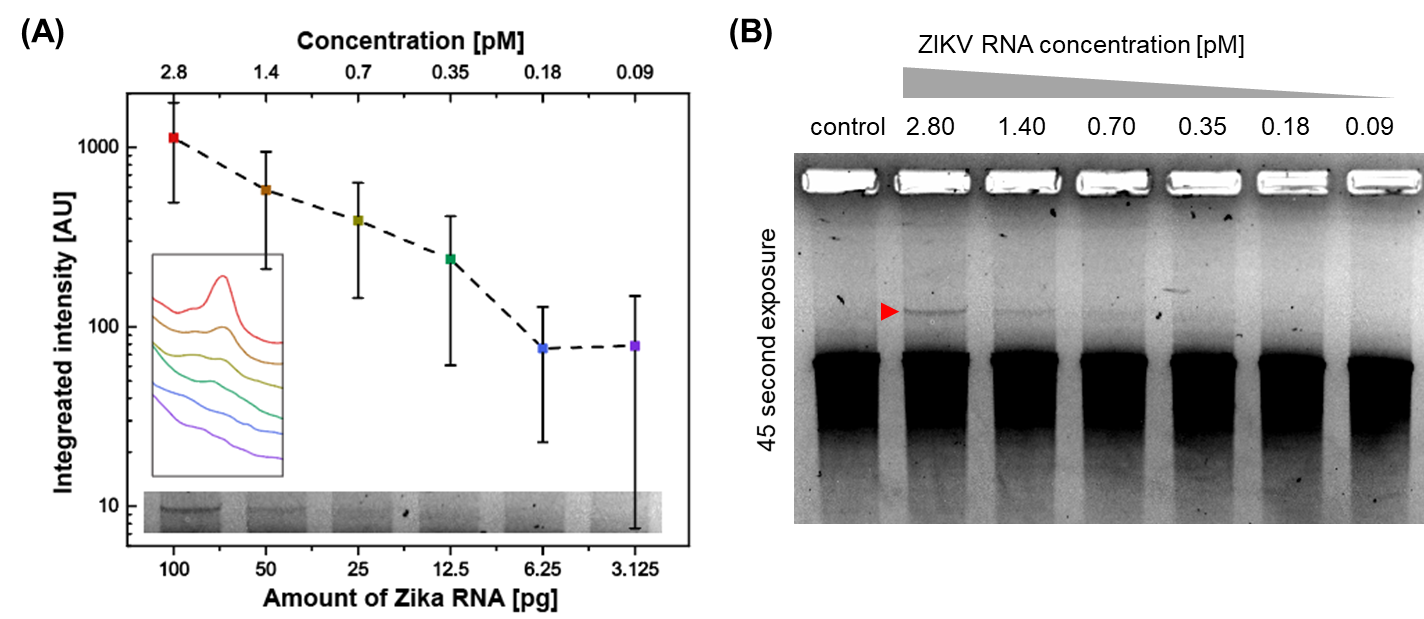

### **Fig. S9. Detection sensitivity test of single nanoswitch**. (**A**) Sensitivity test of a high-performing single nanoswitch (third nanoswitch listed in **Table S3)**. An example of gel image with detection bands is presented at the bottom of the figure and the profiles of the detection bands are presented as an insert figure on the right corner. (**B**) The entire gel image presented at the bottom of (**A**): A visible band can be seen to at least the 1.4 pM lane. Experiment was performed in triplicate and error bars represent the standard deviation.

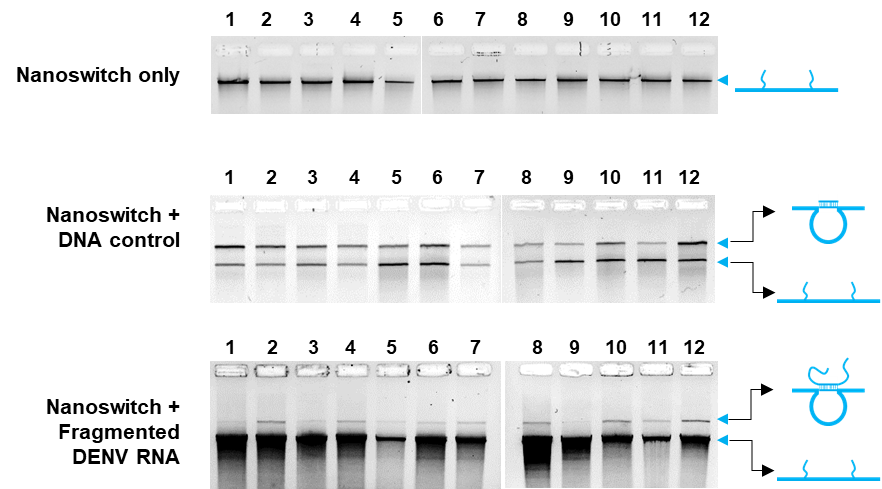

### **Fig. S10. Analysis of the 12 DNA nanoswitches designed for DENV RNA detection.** Top panel shows the negative control test of just the different DNA nanoswitches. The middle panel shows the positive control of complementary ssDNA (2 nM) annealed with the corresponding nanoswitch. The bottom panel shows detection of DENV RNA by individual DNA nanoswitches. We used 10 ng of fragmented *in vitro* transcribed DENV RNA to test the nanoswitches.

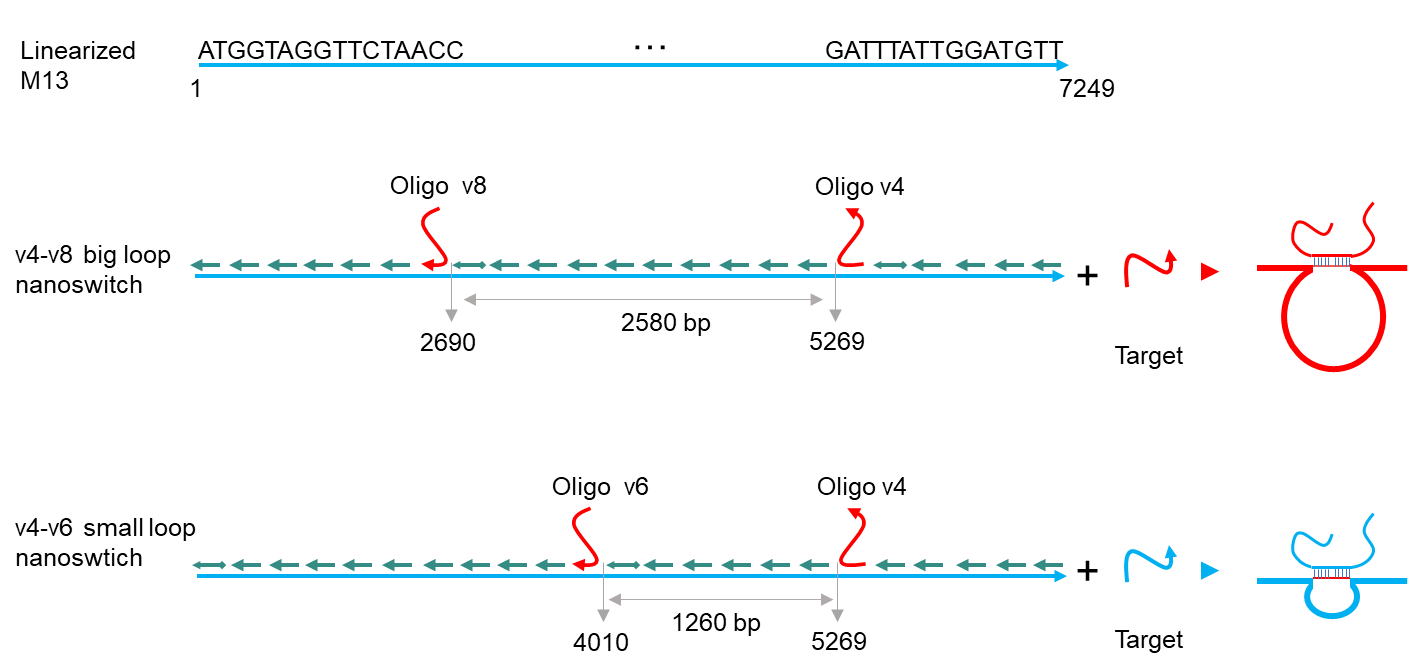

### **Fig. S11. Tuning the loop size of DNA nanoswitch.** The size of v4-v8 loop is about 2580 bp and the size of v4-v6 loop is about 1260 bp. More information can be found in reference ^3,4^. Note in the table of oligos, all detection ssDNA oligos are named with prefix v4- or v8- or v6-.

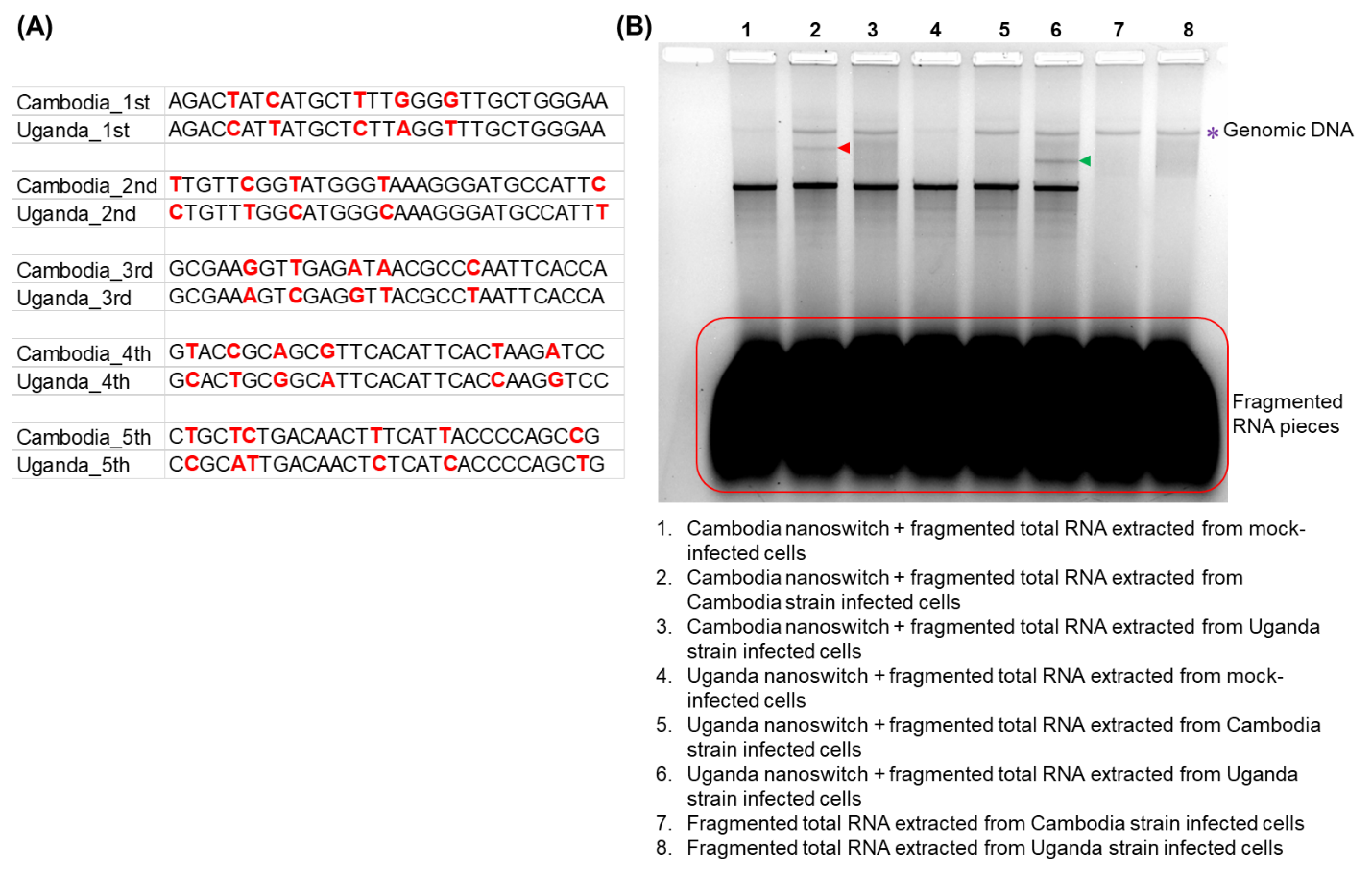

### **Fig. S12. Targets and a gel image of specificity test with Cambodia and Uganda strains of ZIKV.** (**A**) The five targets for the specificity test of Cambodia and Uganda strains of ZIKV. Strain-specific nucleotides are colored in red. (**B**) A representative gel image from the assay demonstrating nanoswitch specificity for detecting and differentiating between ZIKV Cambodia and Uganda strains used in **Fig. 3C-2D** in the main text. * indicates contaminating cellular DNA left in the total RNA and the area in the red frame at the bottom indicates the unbound fragmented pieces of cellular and viral RNA isolated from mock- and ZIKV-infected Huh7 cells. The oligos of corresponding nanoswitches are listed in **Table S7**.

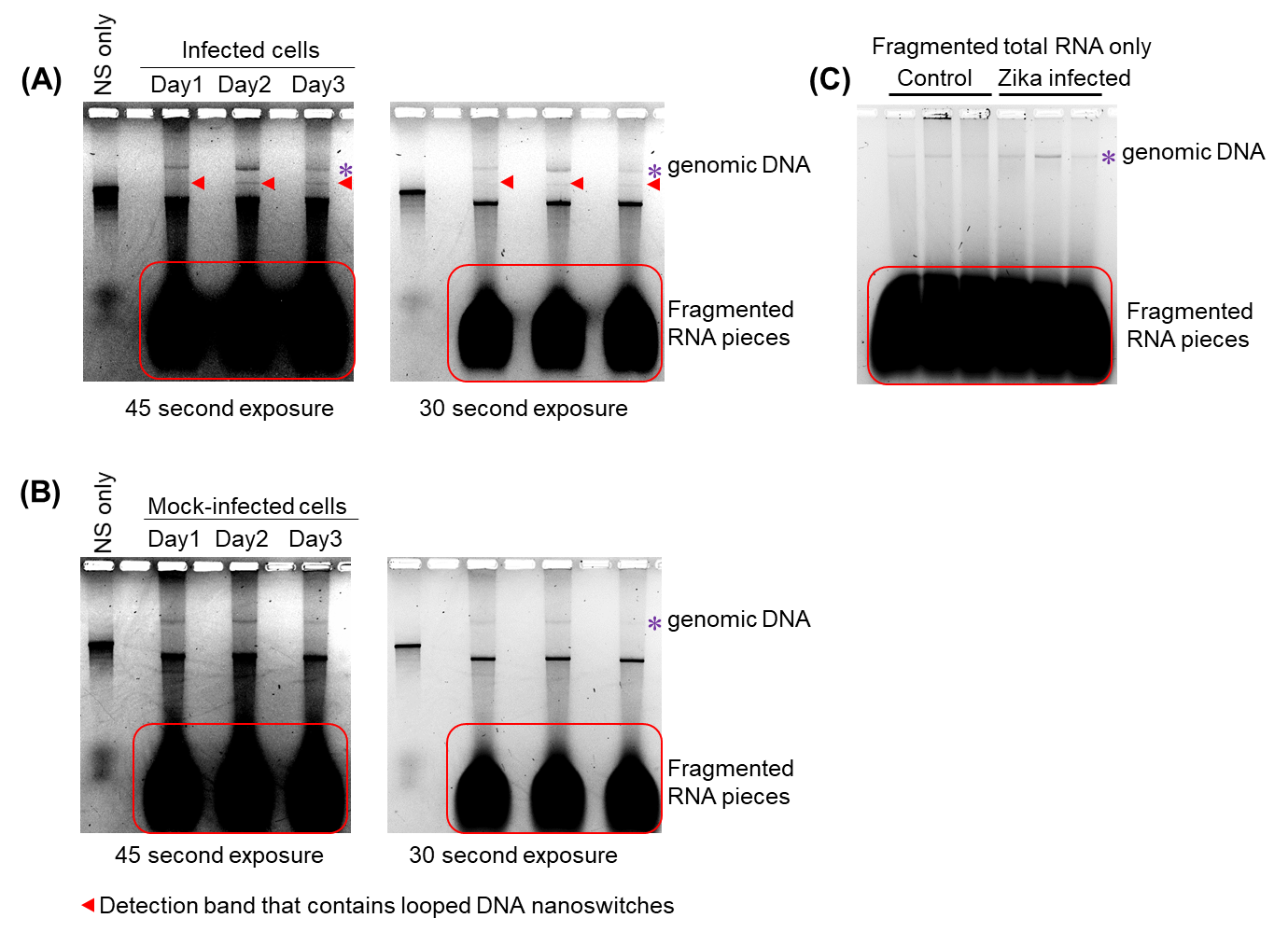

**Fig. S13**. **Detection of ZIKV RNA in total RNA extracted from human liver cells.** (**A**) Detection of ZIKV RNA in total RNA of infected human liver cells, NS: nanoswitch. (**B**) Control experiment used total RNA from mock-infected human liver cells. (**C**) Fragmented total RNA only. Note: the red arrows indicate the detection bands that contain looped DNA nanoswitches and the asterisks indicate the genomic DNA in the total RNA.

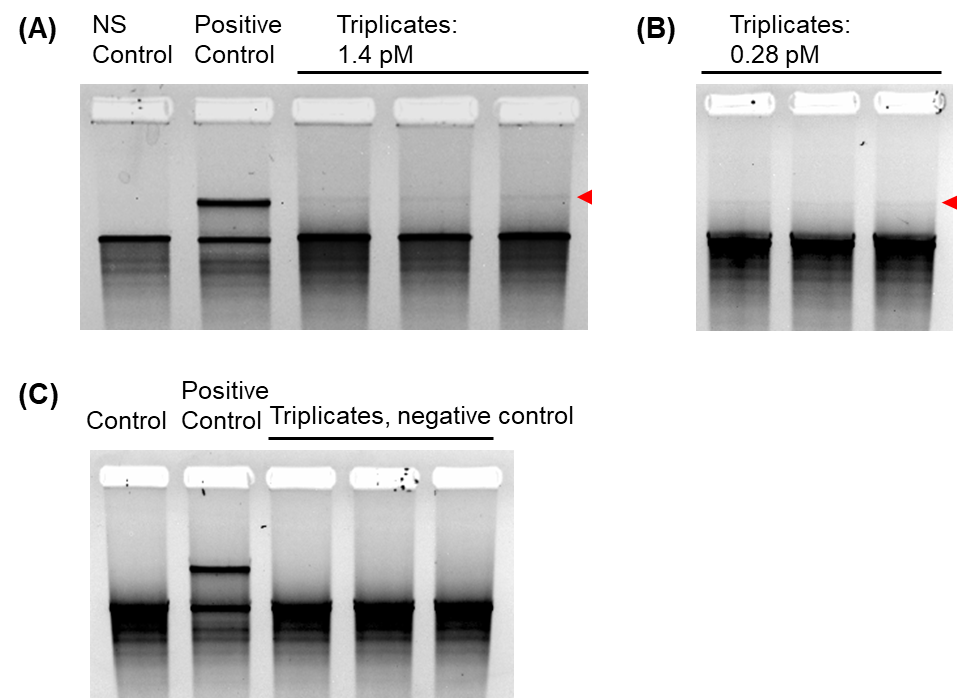

**Fig. S14.** **Gel images of the ZIKV RNA detection in samples mimicking the urine of patients.** Triplicate experiments of detecting ZIKV RNA extracted from human urine at (**A**) 1.4 pM, (**B**) 0.28 pM, and (**C**) 0 pM (negative control). The quantified detection results are presented in **Fig. 5a**.

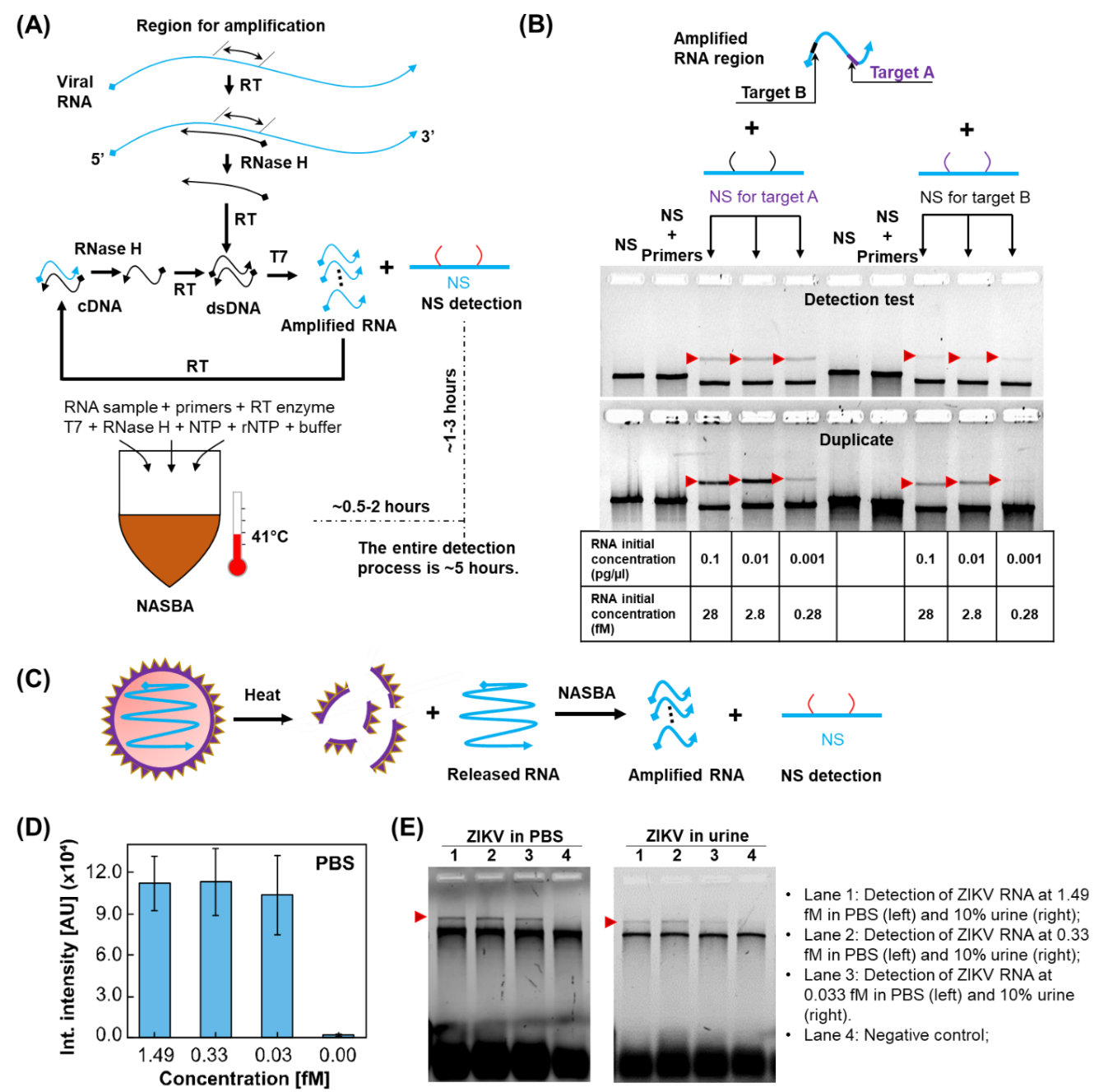

**Fig. S15. Detection of ZIKV RNA based on pre-amplification with NASBA**. (**A**) Basic process of Nucleic Acid Sequence Based Amplification (NASBA), RT: reverse transcription. (**B**) Test of detection based on NASBA amplification. Two targets were chosen on the amplified region of the in vitro transcription ZIKV RNA (targets A and B in **Table S8**). (**C**) Schematic of viral RNA detection based on NASBA. (**D**) Positive detection of ZIKV RNA from infectious virus in PBS. (**E**)Example gel images of the ZIKV RNA detection based on NASBA by spiking virus particles into PBS and urine (final concentration is 10%), the nanoswitch used here is the nanoswitch for target A in **Table S8**.

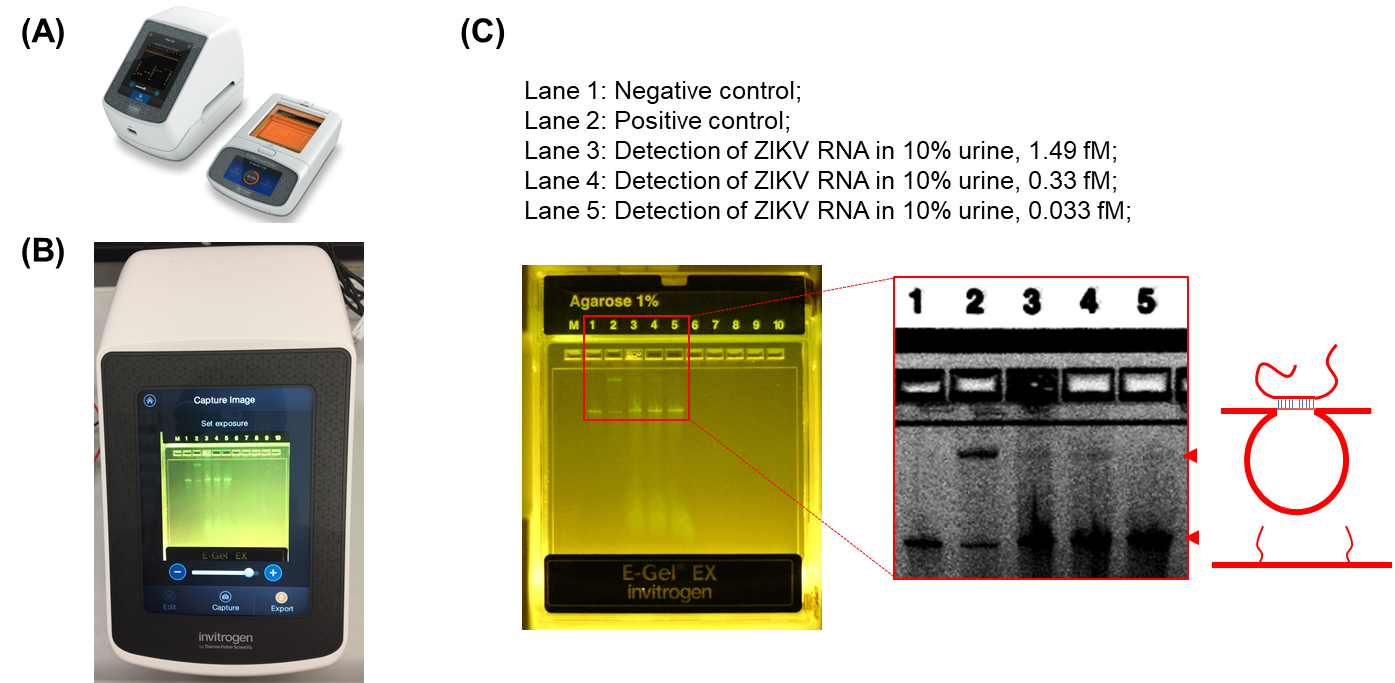

### **Fig. S16. Portable e-gel system for detection of ZIKV RNA based on pre-amplification with NASBA.** (**A**) Commercially available E-gel system, (**B**) Image capture of an E-gel cartridge testing viral nanoswitch detection (run at 48 Volts for 1 hour). (**C**) A gel image of the detection of ZIKV RNA based on pre-amplification with NASBA. The concentrations of ZIKV virus particle in the human urine (10%) are 1.49, 0.33 and 0.033 fM for lane 3, 4 and 5 respectively. The nanoswitch used here is the one for target A in **Table S8**. The E-gel was run at 48 volts for 1 hour.

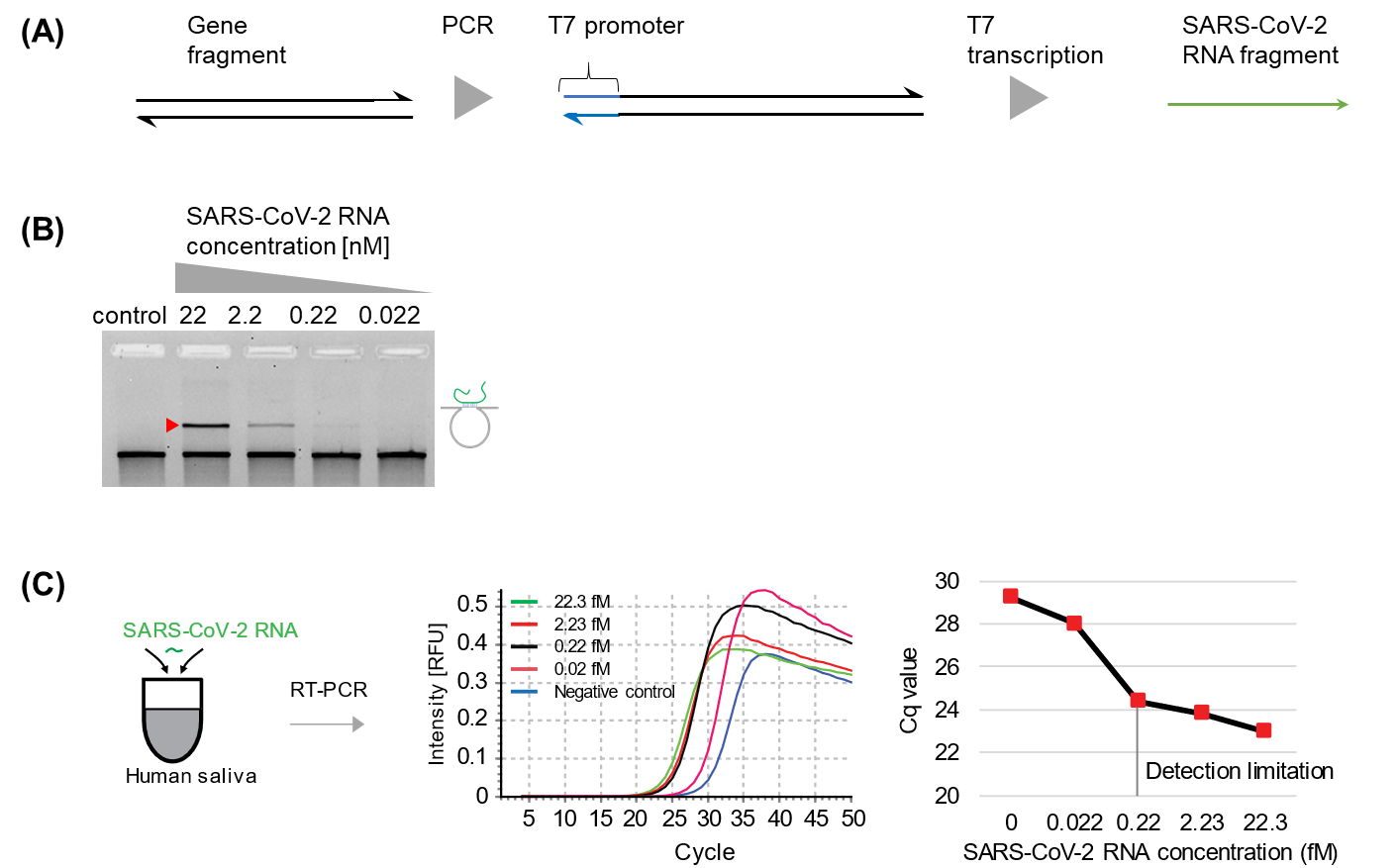

**Figure S17. Detection of SARS-CoV-2 RNA in human saliva**. (**A**) Schematic of producing SARS-CoV-2 RNA fragment. (**B**) Detection test of SARS-CoV-2 RNA with different concentration in buffer. (**C**) RT-PCR detection of SARS-CoV-2 RNA in 10% human saliva. Based on the Cq value shown on the right, the detection limitation of RT-PCR in this scenario is about 0.22 fM.

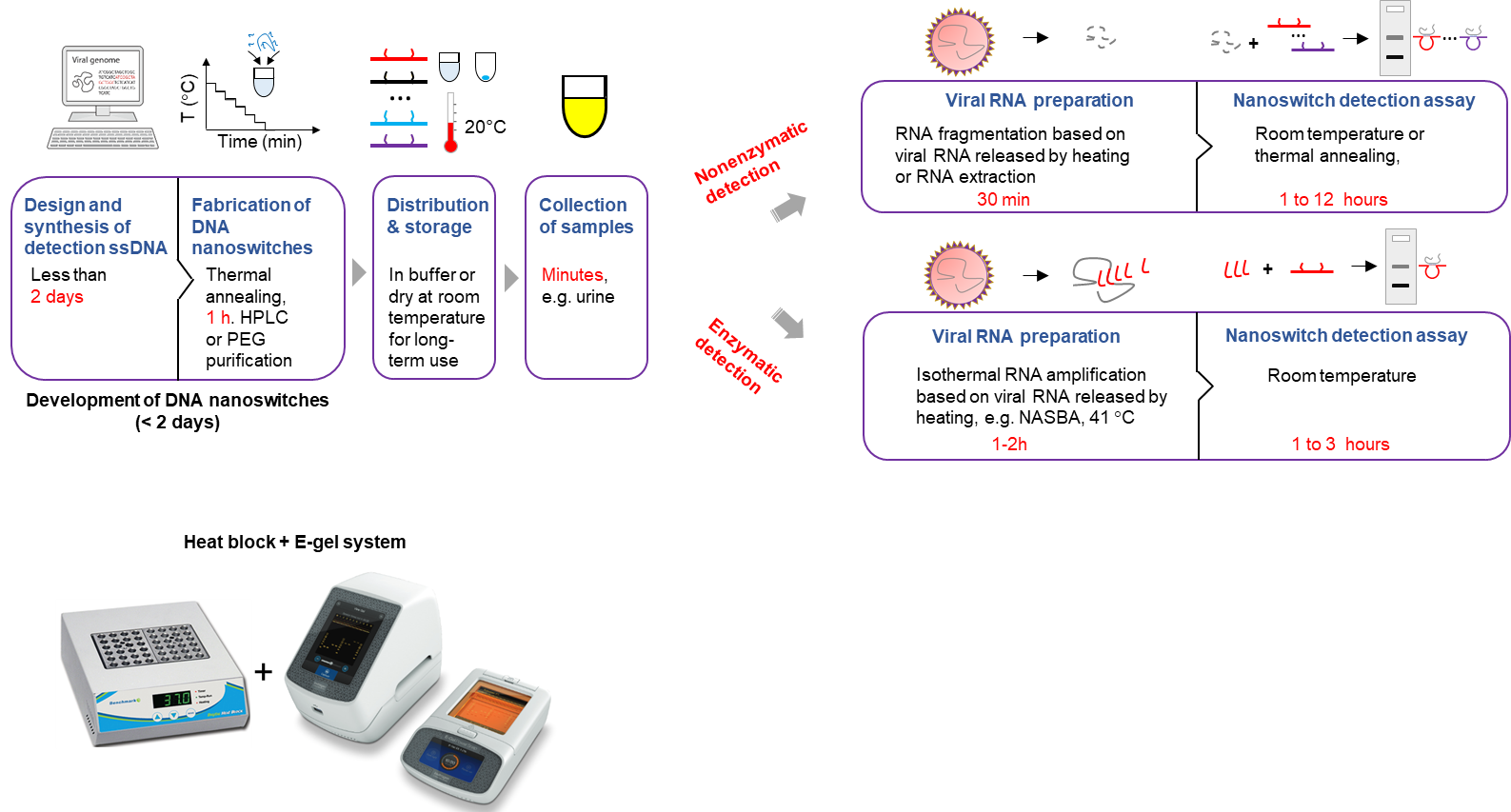

**Fig. S18. Development cycle for DNA nanoswitch based detection of viral RNAs.** Direct detection can be accomplished in ~1-13 hours and in only 2-5 hours with pre-amplification. Bottom left shows the minimum equipment (heat block and E-gel system, pipettes, tips, and tubes are not shown here) needed for our method.

### **Table S1.** A ZIKV RNA target sequence from a literature^5^ and its corresponding detector ssDNA (**Fig. 2c**).

| **Name** | **Sequence (5’-3’)** | **Length** |
| --- | --- | --- |
| ZIKV_s1_Target | AGCCTACCTTGACAAGCAATCAGACACTCA | 30 |
| v4-ZIKV_s1_40-15 | ACCGTTGTAGCAATACTTCTTTGATTAGTAATAACATCAC**TGAGTGTCTGATTGC** | 55 |
| v8-ZIKV_s1_ 15-40 | **TTGTCAAGGTAGGCT**TCAACCGATTGAGGGAGGGAAGGTAAATATTGACGGAAAT | 55 |

### **Table S2.** Target sequence and different lengths of detector ssDNA (15, 14, 13, 12, 11, 10nt) for optimization the design of nanoswitch (**Fig. S3**).

| **Name** | **Sequence (5’-3’)** | **Length** |
| --- | --- | --- |
| ZIKV_arm length test_Target | AACGCCCAATTCACCAAGAGCCGAAGCCAC | 30 |
| v4-ZIKV arm length test 30-15 | CAATACTTCTTTGATTAGTAATAACATCAC***GTGGCTTCGGCTCTT*** | 45 |
| v8-ZIKV arm length test 15-30 | ***GGTGAATTGGGCGTT***TCAACCGATTGAGGGAGGGAAGGTAAATAT | 45 |
| v4-ZIKV arm length test 30-14 | CAATACTTCTTTGATTAGTAATAACATCAC***TGGCTTCGGCTCTT*** | 44 |
| v8-ZIKV arm length test 14-30 | ***GGTGAATTGGGCGT***TCAACCGATTGAGGGAGGGAAGGTAAATAT | 44 |
| v4-ZIKV arm length test 30-13 | CAATACTTCTTTGATTAGTAATAACATCAC***GGCTTCGGCTCTT*** | 43 |
| v8-ZIKV arm length test 13-30 | ***GGTGAATTGGGCG***TCAACCGATTGAGGGAGGGAAGGTAAATAT | 43 |
| v4-ZIKV arm length test 30-12 | CAATACTTCTTTGATTAGTAATAACATCAC***GCTTCGGCTCTT*** | 42 |
| v8-ZIKV arm length test 12-30 | ***GGTGAATTGGGC***TCAACCGATTGAGGGAGGGAAGGTAAATAT | 42 |
| v4-ZIKV arm length test 30-11 | CAATACTTCTTTGATTAGTAATAACATCAC***CTTCGGCTCTT*** | 41 |
| v8-ZIKV arm length test 11-30 | ***GGTGAATTGGG***TCAACCGATTGAGGGAGGGAAGGTAAATAT | 41 |
| v4-ZIKV arm length test 30-10 | CAATACTTCTTTGATTAGTAATAACATCAC***TTCGGCTCTT*** | 40 |
| v8-ZIKV arm length test 10-30 | ***GGTGAATTGG***TCAACCGATTGAGGGAGGGAAGGTAAATAT | 40 |

### **Table S3**. The eighteen target sequences and corresponding detector ssDNA oligos for the detection of ZIKV RNA (**Fig. 2e, 2f, 3a, 3b, 3d, 5a and S7, S8, S9**).

Note: the position of each target sequence on the ZIKV RNA is shown in the far-right column.

| **Nanoswitch** | **Name** | **Sequence (5’-3’)** | **Len.** | **Pos.** |
| --- | --- | --- | --- | --- |
| 1 | ZIKV_Target1 | GTGTGATGCCACCATGAGCTATGAATGCCC | 30 | 605 |
|  | v4-ZIKV T1 40-15 | ACCGTTGTAGCAATACTTCTTTGATTAGTAATAACATCAC**GGGCATTCATAGCTC** | 55 |  |
|  | v8-ZIKV T1 15-40 | **ATGGTGGCATCACAC**TCAACCGATTGAGGGAGGGAAGGTAAATATTGACGGAAAT | 55 |  |
| 2 | ZIKV_Target2 | AGTGGACAGAGGCTGGGGAAATGGATGTGG | 30 | 1265 |
|  | v4-ZIKV T2 40-15 | ACCGTTGTAGCAATACTTCTTTGATTAGTAATAACATCAC**CCACATCCATTTCCC** | 55 |  |
|  | v8-ZIKV T2 15-40 | **CAGCCTCTGTCCACT**TCAACCGATTGAGGGAGGGAAGGTAAATATTGACGGAAAT | 55 |  |
| 3 | ZIKV_Target3 | AACGCCCAATTCACCAAGAGCCGAAGCCAC | 30 | 1484 |
|  | v4-ZIKV T3 40-15 | ACCGTTGTAGCAATACTTCTTTGATTAGTAATAACATCAC**GTGGCTTCGGCTCTT** | 55 |  |
|  | v8-ZIKV T3 15-40 | **GGTGAATTGGGCGTT**TCAACCGATTGAGGGAGGGAAGGTAAATATTGACGGAAAT | 55 |  |
| 4 | ZIKV_Target4 | AGGGAGTCAAGAAGGAGCAGTTCACACGGC | 30 | 1751 |
|  | v4-ZIKV T4 40-15 | ACCGTTGTAGCAATACTTCTTTGATTAGTAATAACATCAC**GCCGTGTGAACTGCT** | 55 |  |
|  | v8-ZIKV T4 15-40 | **CCTTCTTGACTCCCT**TCAACCGATTGAGGGAGGGAAGGTAAATATTGACGGAAAT | 55 |  |
| 5 | ZIKV_Target5 | GTACCATCCTGACTCCCCTCGTAGATTGGC | 30 | 2588 |
|  | v4-ZIKV T5 40-15 | ACCGTTGTAGCAATACTTCTTTGATTAGTAATAACATCAC**GCCAATCTACGAGGG** | 55 |  |
|  | v8-ZIKV T5 15-40 | **GAGTCAGGATGGTAC**TCAACCGATTGAGGGAGGGAAGGTAAATATTGACGGAAAT | 55 |  |
| 6 | ZIKV_Target6 | ACATCATGTGGAGATCAGTAGAAGGGGAGC | 30 | 2683 |
|  | v4-ZIKV T6 40-15 | ACCGTTGTAGCAATACTTCTTTGATTAGTAATAACATCAC**GCTCCCCTTCTACTG** | 55 |  |
|  | v8-ZIKV T6 15-40 | **ATCTCCACATGATGT**TCAACCGATTGAGGGAGGGAAGGTAAATATTGACGGAAAT | 55 |  |
| 7 | ZIKV_Target7 | GAAGAACGACACATGGAGGCTGAAGAGGGC | 30 | 3104 |
|  | v4-ZIKV T7 40-15 | ACCGTTGTAGCAATACTTCTTTGATTAGTAATAACATCAC**GCCCTCTTCAGCCTC** | 55 |  |
|  | v8-ZIKV T7 15-40 | **CATGTGTCGTTCTTC**TCAACCGATTGAGGGAGGGAAGGTAAATATTGACGGAAAT | 55 |  |
| 8 | ZIKV Target 8 | CTAATTGGACACCCCGTGAGAGCATGCTGC | 30 | 3835 |
|  | v4-ZIKV T8 40-15 | ACCGTTGTAGCAATACTTCTTTGATTAGTAATAACATCAC**GCAGCATGCTCTCAC** | 55 |  |
|  | v8-ZIKV T8 15-40 | **GGGGTGTCCAATTAG**TCAACCGATTGAGGGAGGGAAGGTAAATATTGACGGAAAT | 55 |  |
| 9 | ZIKV_Target9 | AAACAGTCCCCGGCTCGATGTGGCACTAGA | 30 | 4430 |
|  | v4-ZIKV T9 40-15 | ACCGTTGTAGCAATACTTCTTTGATTAGTAATAACATCAC**TCTAGTGCCACATCG** | 55 |  |
|  | v8-ZIKV T9 15-40 | **AGCCGGGGACTGTTT**TCAACCGATTGAGGGAGGGAAGGTAAATATTGACGGAAAT | 55 |  |
| 10 | ZIKV_Target10 | CCCGGAGAGAGAGCGAGGAACATCCAGACT | 30 | 4917 |
|  | v4-ZIKV T10 40-15 | ACCGTTGTAGCAATACTTCTTTGATTAGTAATAACATCAC**AGTCTGGATGTTCCT** | 55 |  |
|  | v8-ZIKV T10 15-40 | **CGCTCTCTCTCCGGG**TCAACCGATTGAGGGAGGGAAGGTAAATATTGACGGAAAT | 55 |  |
| 11 | ZIKV_Target11 | GGACTACCCAGCAGGAACTTCAGGATCTCC | 30 | 4997 |
|  | v4-ZIKV T11 40-15 | ACCGTTGTAGCAATACTTCTTTGATTAGTAATAACATCAC**GGAGATCCTGAAGTT** | 55 |  |
|  | v8-ZIKV T11 15-40 | **CCTGCTGGGTAGTCC**TCAACCGATTGAGGGAGGGAAGGTAAATATTGACGGAAAT | 55 |  |
| 12 | ZIKV_Target12 | GTGACGCATTCCCGGACTCCAACTCACCAA | 30 | 5581 |
|  | v4-ZIKV T12 40-15 | ACCGTTGTAGCAATACTTCTTTGATTAGTAATAACATCAC**TTGGTGAGTTGGAGT** | 55 |  |
|  | v8-ZIKV T12 15-40 | **CCGGGAATGCGTCAC**TCAACCGATTGAGGGAGGGAAGGTAAATATTGACGGAAAT | 55 |  |
| 13 | ZIKV_Target13 | GAGTTCCAGAAAACAAAACATCAAGAGTGG | 30 | 5793 |
|  | v4-ZIKV T13 40-15 | ACCGTTGTAGCAATACTTCTTTGATTAGTAATAACATCAC**CCACTCTTGATGTTT** | 55 |  |
|  | v8-ZIKV T13 15-40 | **TGTTTTCTGGAACTC**TCAACCGATTGAGGGAGGGAAGGTAAATATTGACGGAAAT | 55 |  |
| 14 | ZIKV_Target14 | CATCTAATGGGAAGGAGAGAGGAGGGGGCA | 30 | 6957 |
|  | v4-ZIKV T14 40-15 | ACCGTTGTAGCAATACTTCTTTGATTAGTAATAACATCAC**TGCCCCCTCCTCTCT** | 55 |  |
|  | v8-ZIKV T14 15-40 | **CCTTCCCATTAGATG**TCAACCGATTGAGGGAGGGAAGGTAAATATTGACGGAAAT | 55 |  |
| 15 | ZIKV_Target15 | CACAGGAATAGCCATGACCGACACCACACC | 30 | 8684 |
|  | v4-ZIKV T15 40-15 | ACCGTTGTAGCAATACTTCTTTGATTAGTAATAACATCAC**GGTGTGGTGTCGGTC** | 55 |  |
|  | v8-ZIKV T15 15-40 | **ATGGCTATTCCTGTG**TCAACCGATTGAGGGAGGGAAGGTAAATATTGACGGAAAT | 55 |  |
| 16 | ZIKV_Target16 | GGATGGGGAGAGAGAATTCAGGAGGTGGTG | 30 | 9160 |
|  | v4-ZIKV T16 40-15 | ACCGTTGTAGCAATACTTCTTTGATTAGTAATAACATCAC**CACCACCTCCTGAAT** | 55 |  |
|  | v8-ZIKV T16 15-40 | **TCTCTCTCCCCATCC**TCAACCGATTGAGGGAGGGAAGGTAAATATTGACGGAAAT | 55 |  |
| 17 | ZIKV_Target17 | GAGGAAGTTCTAGAGATGCAAGACTTGTGG | 30 | 9549 |
|  | v4-ZIKV T17 40-15 | ACCGTTGTAGCAATACTTCTTTGATTAGTAATAACATCAC**CCACAAGTCTTGCAT** | 55 |  |
|  | v8-ZIKV T17 15-40 | **CTCTAGAACTTCCTC**TCAACCGATTGAGGGAGGGAAGGTAAATATTGACGGAAAT | 55 |  |
| 18 | ZIKV_Target18 | CTGAGTCAAAAAACCCCACGCGCTTGGAGG | 30 | 10543 |
|  | v4-ZIKV T18 40-15 | ACCGTTGTAGCAATACTTCTTTGATTAGTAATAACATCAC**CCTCCAAGCGCGTGG** | 55 |  |
|  | v8-ZIKV T18 15-40 | **GGTTTTTTGACTCAG**TCAACCGATTGAGGGAGGGAAGGTAAATATTGACGGAAAT | 55 |  |

### **Table S4**. The twelve target sequences and corresponding detector ssDNA oligos for the detection of DENV RNA (**Fig. 3a, S10**).

Note: the position of each target sequence on the DENV RNA is shown in the far-right column.

| **Nanoswitch** | **Name** | **Sequence (5’-3’)** | **Len.** | **Pos.** |
| --- | --- | --- | --- | --- |
| 1 | DENV Target 1 | GTGACTGAGGACTGCGGAAATAGAGGACCC | 30 | 2823 |
|  | v4-DENV T1 40-15 | ACCGTTGTAGCAATACTTCTTTGATTAGTAATAACATCAC**GGGTCCTCTATTTCC** | 55 |  |
|  | v8-DENV T1 15-40 | **GCAGTCCTCAGTCAC**TCAACCGATTGAGGGAGGGAAGGTAAATATTGACGGAAAT | 55 |  |
| 2 | DENV Target 2 | CTCTCCTCCCAGAGCACTATACCAGAGACC | 30 | 3280 |
|  | v4-DENV T2 40-15 | ACCGTTGTAGCAATACTTCTTTGATTAGTAATAACATCAC**GGTCTCTGGTATAGT** | 55 |  |
|  | v8-DENV T2 15-40 | **GCTCTGGGAGGAGAG**TCAACCGATTGAGGGAGGGAAGGTAAATATTGACGGAAAT | 55 |  |
| 3 | DENV Target 3 | TGCTCACTGGACGATCGGCCGATTTGGAAC | 30 | 3805 |
|  | v4-DENV T3 40-15 | ACCGTTGTAGCAATACTTCTTTGATTAGTAATAACATCAC**GTTCCAAATCGGCCG** | 55 |  |
|  | v8-DENV T3 15-40 | **ATCGTCCAGTGAGCA**TCAACCGATTGAGGGAGGGAAGGTAAATATTGACGGAAAT | 55 |  |
| 4 | DENV Target 4 | GGCCAGCACTCCAAGCAAAAGCATCCAGAG | 30 | 4259 |
|  | v4-DENV T4 40-15 | ACCGTTGTAGCAATACTTCTTTGATTAGTAATAACATCAC**CTCTGGATGCTTTTG** | 55 |  |
|  | v8-DENV T4 15-40 | **CTTGGAGTGCTGGCC**TCAACCGATTGAGGGAGGGAAGGTAAATATTGACGGAAAT | 55 |  |
| 5 | DENV Target 5 | CACACCAGAAGGGAAAGTAGTGGACCTCGG | 30 | 5488 |
|  | v4-DENV T5 40-15 | ACCGTTGTAGCAATACTTCTTTGATTAGTAATAACATCAC**CCGAGGTCCACTACT** | 55 |  |
|  | v8-DENV T5 15-40 | **TTCCCTTCTGGTGTG**TCAACCGATTGAGGGAGGGAAGGTAAATATTGACGGAAAT | 55 |  |
| 6 | DENV Target 6 | AAGCCACTTACGAGCCGGATGTTGACCTCG | 30 | 6432 |
|  | v4-DENV T6 40-15 | ACCGTTGTAGCAATACTTCTTTGATTAGTAATAACATCAC**CGAGGTCAACATCCG** | 55 |  |
|  | v8-DENV T6 15-40 | **GCTCGTAAGTGGCTT**TCAACCGATTGAGGGAGGGAAGGTAAATATTGACGGAAAT | 55 |  |
| 7 | DENV Target 7 | GCATGGCGTAGTGGACTAGCGGTTAGAGGA | 30 | 7190 |
|  | v4-DENV T7 40-15 | ACCGTTGTAGCAATACTTCTTTGATTAGTAATAACATCAC**TCCTCTAACCGCTAG** | 55 |  |
|  | v8-DENV T7 15-40 | **TCCACTACGCCATGC**TCAACCGATTGAGGGAGGGAAGGTAAATATTGACGGAAAT | 55 |  |
| 8 | DENV Target 8 | CAAGCTACAGCTCAAAGGAATGTCATACTC | 30 | 7782 |
|  | v4-DENV T8 40-15 | ACCGTTGTAGCAATACTTCTTTGATTAGTAATAACATCAC**GAGTATGACATTCCT** | 55 |  |
|  | v8-DENV T8 15-40 | **TTGAGCTGTAGCTTG**TCAACCGATTGAGGGAGGGAAGGTAAATATTGACGGAAAT | 55 |  |
| 9 | DENV Target 9 | GACCCATTTCCTCAGAGCAATGCACCAATC | 30 | 8315 |
|  | v4-DENV T9 40-15 | ACCGTTGTAGCAATACTTCTTTGATTAGTAATAACATCAC**GATTGGTGCATTGCT** | 55 |  |
|  | v8-DENV T9 15-40 | **CTGAGGAAATGGGTC**TCAACCGATTGAGGGAGGGAAGGTAAATATTGACGGAAAT | 55 |  |
| 10 | DENV Target 13 | GAAGGCAAGAAACGCACTGGACAACTTAGC | 30 | 8653 |
|  | v4-DENV T10 40-15 | ACCGTTGTAGCAATACTTCTTTGATTAGTAATAACATCAC**GCTAAGTTGTCCAGT** | 55 |  |
|  | v8-DENV T10 15-40 | **GCGTTTCTTGCCTTC**TCAACCGATTGAGGGAGGGAAGGTAAATATTGACGGAAAT | 55 |  |
| 11 | DENV Target 11 | AGAACCCAAGAACCGAAAGAAGGCACGAAG | 30 | 9313 |
|  | v4-DENV T11 40-15 | ACCGTTGTAGCAATACTTCTTTGATTAGTAATAACATCAC**CTTCGTGCCTTCTTT** | 55 |  |
|  | v8-DENV T11 15-40 | **CGGTTCTTGGGTTCT**TCAACCGATTGAGGGAGGGAAGGTAAATATTGACGGAAAT | 55 |  |
| 12 | DENV Target 12 | AGACCAACACCAAGAGGCACAGTAATGGAC | 30 | 10481 |
|  | v4-DENV T12 40-15 | ACCGTTGTAGCAATACTTCTTTGATTAGTAATAACATCAC**GTCCATTACTGTGCC** | 55 |  |
|  | v8-DENV T12 15-40 | **TCTTGGTGTTGGTCT**TCAACCGATTGAGGGAGGGAAGGTAAATATTGACGGAAAT | 55 |  |

### **Table S5.** Variable oligos for constructing nanoswitches with different loop sizes used in **Fig. 3b, 3d**.

|  | **Name** | **Sequence (5’-3’)** | **Length** |
| --- | --- | --- | --- |
| **For v4-v8 loop nanoswitch** | v4 oligo | Oligos with prefix ‘v4-’ |  |
|  | v8 oligo | Oligos with prefix ‘v8-’ |  |
|  | Var 4 filler | TCTGTCCATCACGCAAATTA | 20 |
|  | Var 8 filler | TATTCATTAAAGGTGAATTA | 20 |
| **For v4-v6 loop nanoswitch** | v4 oligo | Oligos with prefix ‘v4-’ |  |
|  | v8 oligo | Oligos with prefix ‘v6-’ |  |
|  | Var 4 filler | TCTGTCCATCACGCAAATTA | 20 |
|  | Var 6 filler | TCGCAAGACAAAGAACGCGA | 20 |
| **For v4-v7 loop nanoswitch** | v4 oligo | Oligos with prefix ‘v4-’ |  |
|  | v7 oligo | Oligos with prefix ‘v7-’ | 20 |
|  | Var 4 filler | TCTGTCCATCACGCAAATTA | 20 |
|  | Var 7 filler | TCGCAAGACAAAGAACGCGA |  |

### **Table S6.** Target sequences and the corresponding detector ssDNA for the ZIKV and DENV multiplexing test (**Fig. 3b**).

| **Name** | **Sequence (5’-3’)** | **Len.** |
| --- | --- | --- |
| ZIKV_Target3 | AACGCCCAATTCACCAAGAGCCGAAGCCAC | 30 |
| v4-ZIKV T3 40-15 | ACCGTTGTAGCAATACTTCTTTGATTAGTAATAACATCAC**GTGGCTTCGGCTCTT** | 55 |
| v8-ZIKV T3 15-40 | **GGTGAATTGGGCGTT**TCAACCGATTGAGGGAGGGAAGGTAAATATTGACGGAAAT | 55 |
| DENV_Target 10 | GCATGGCGTAGTGGACTAGCGGTTAGAGGA | 30 |
| v4-DENV T10 40-15 | ACCGTTGTAGCAATACTTCTTTGATTAGTAATAACATCAC**TCCTCTAACCGCTAG** | 55 |
| v6-DENV T10 15-40 | **TCCACTACGCCATGC**TGGGTTATATAACTATATGTAAATGCTGATGCAAATCCAA | 55 |

### **Table S7.** Target sequences and the corresponding detection arm ssDNA for the ZIKV Cambodia and Uganda specificity test (**Fig. 3d**).

| **Name** | **Sequence (5’-3’)** | **Len.** |
| --- | --- | --- |
| Cambodia_1st | AGAC**T**AT**C**ATGCT**T**TT**G**GG**G**TTGCTGGGAA | 30 |
| v4-Cambodia_1st_ 40_15 | ACCGTTGTAGCAATACTTCTTTGATTAGTAATAACATCAC**TTCCCAGCAACCCCA** | 55 |
| v8-Cambodia_1st_15_40 | **AAAGCATGATAGTCT**TCAACCGATTGAGGGAGGGAAGGTAAATATTGACGGAAAT | 55 |
| Cambodia_2nd | **T**TGTT**C**GG**T**ATGGG**T**AAAGGGATGCCATT**C** | 30 |
| v4-Cambodia_2nd_ 40_15 | ACCGTTGTAGCAATACTTCTTTGATTAGTAATAACATCAC**GAATGGCATCCCTTT** | 55 |
| v8-Cambodia_2nd_ 15_40 | **ACCCATACCGAACAA**TCAACCGATTGAGGGAGGGAAGGTAAATATTGACGGAAAT | 55 |
| Cambodia_3rd | GCGAA**G**GT**T**GAG**A**T**A**ACGCC**C**AATTCACCA | 30 |
| v4-Cambodia_3rd_ 40_15 | ACCGTTGTAGCAATACTTCTTTGATTAGTAATAACATCAC**TGGTGAATTGGGCGT** | 55 |
| v8-Cambodia_3rd_ 15_40 | **TATCTCAACCTTCGC**TCAACCGATTGAGGGAGGGAAGGTAAATATTGACGGAAAT | 55 |
| Cambodia_4th | G**T**AC**C**GC**A**GC**G**TTCACATTCAC**T**AAG**A**TCC | 30 |
| v4-Cambodia_4th_ 40_15 | ACCGTTGTAGCAATACTTCTTTGATTAGTAATAACATCAC**GGATCTTAGTGAATG** | 55 |
| v8-Cambodia_4th_ 15_40 | **TGAACGCTGCGGTAC**TCAACCGATTGAGGGAGGGAAGGTAAATATTGACGGAAAT | 55 |
| Cambodia_5th | C**T**GC**TC**TGACAACT**T**TCAT**T**ACCCCAGC**C**G | 30 |
| v4-Cambodia_5th_ 40_15 | ACCGTTGTAGCAATACTTCTTTGATTAGTAATAACATCAC**CGGCTGGGGTAATGA** | 55 |
| v8-Cambodia_5th_ 15_40 | **AAGTTGTCAGAGCAG**TCAACCGATTGAGGGAGGGAAGGTAAATATTGACGGAAAT | 55 |
| Uganda_1st | AGAC**C**AT**T**ATGCT**C**TT**A**GG**T**TTGCTGGGAA | 30 |
| v4-Uganda_1st_40_15 | ACCGTTGTAGCAATACTTCTTTGATTAGTAATAACATCAC**TTCCCAGCAAACCTA** | 55 |
| v7-Uganda_1st_15_40 | **AGAGCATAATGGTCT**GTTTTAGCGAACCTCCCGACTTGCGGGAGGTTTTGAAGCC | 55 |
| Uganda_2nd | **C**TGTT**T**GG**C**ATGGG**C**AAAGGGATGCCATT**T** | 30 |
| v4-Uganda_2nd_40_15 | ACCGTTGTAGCAATACTTCTTTGATTAGTAATAACATCAC**AAATGGCATCCCTTT** | 55 |
| v7-Uganda_2nd_15_40 | **GCCCATGCCAAACAG**GTTTTAGCGAACCTCCCGACTTGCGGGAGGTTTTGAAGCC | 55 |
| Uganda_3rd | GCGAA**A**GT**C**GAG**G**T**T**ACGCC**T**AATTCACCA | 30 |
| v4-Uganda_3rd_40_15 | ACCGTTGTAGCAATACTTCTTTGATTAGTAATAACATCAC**TGGTGAATTAGGCGT** | 55 |
| v7-Uganda_3rd_15_40 | **AACCTCGACTTTCGC**GTTTTAGCGAACCTCCCGACTTGCGGGAGGTTTTGAAGCC | 55 |
| Uganda_4th | G**C**AC**T**GC**G**GC**A**TTCACATTCAC**C**AAG**G**TCC | 30 |
| v4-Uganda_4th_40_15 | ACCGTTGTAGCAATACTTCTTTGATTAGTAATAACATCAC**GGACCTTGGTGAATG** | 55 |
| v7-Uganda_4th_15_40 | **TGAATGCCGCAGTGC**GTTTTAGCGAACCTCCCGACTTGCGGGAGGTTTTGAAGCC | 55 |
| Uganda_5th | C**C**GC**AT**TGACAACT**C**TCAT**C**ACCCCAGC**T**G | 30 |
| v4-Uganda_5th_40_15 | ACCGTTGTAGCAATACTTCTTTGATTAGTAATAACATCAC**CAGCTGGGGTGATGA** | 55 |
| v7-Uganda_5th_15_40 | **GAGTTGTCAATGCGG**GTTTTAGCGAACCTCCCGACTTGCGGGAGGTTTTGAAGCC | 55 |

### **Table S8.** Amplified region of ZIKV RNA, primers, targets and corresponding detector ssDNA used in NASBA related experiments in **Fig. 5b, S15, S16**. The region is chosen based on the reference^6^.

The forward primer has a T7 promoter: AATTCTAATACGACTCACTATAGGGAGAAGG.

| **Name** | **Sequence (5’-3’)** | **Len.** |
| --- | --- | --- |
| Amplified region on ZIKV RNA (1394-1560) | **AATGCTGTCAGTTCATGGCTCCCA**GCACAGTGGGATGATCGTTAATGATACAGGACATGAAACTGATGAGAATAGAGCGAAGGTTGAGATAACGCCCAATTCACCAAGAGCCGAAGCCACCCTGGGGGGTTTTGGAAGCCTAGGACT**TGATTGTGAACCGAGGACAG** | 167 |
| ZIKV NASBA_Reverse primer | CTGTCCTCGGTTCACAATCA | 20 |
| ZIKV NASBA_Forward primer | **AATTCTAATACGACTCACTATAGGGAGAAGG**AATGCTGTCAGTTCATGGCTCCCA | 55 |
| ZIKV_NASBA_Target A | AACGCCCAATTCACCAAGAGCCGAAGCCAC | 30 |
| v4-ZIKV NASBA_Target A 40-15 | ACCGTTGTAGCAATACTTCTTTGATTAGTAATAACATCAC**GTGGCTTCGGCTCTT** | 55 |
| v8-ZIKV NASBA_Target A 15-40 | **GGTGAATTGGGCGTT**TCAACCGATTGAGGGAGGGAAGGTAAATATTGACGGAAAT | 55 |
| ZIKV_NASBA_Target B | GATGATCGTTAATGATACAGGACATGAAAC | 30 |
| v4-ZIKV NASBA_Target B 40-15 | ACCGTTGTAGCAATACTTCTTTGATTAGTAATAACATCAC**GTTTCATGTCCTGTA** | 55 |
| v8-ZIKV NASBA_Target B 15-40 | **TCATTAACGATCATC**TCAACCGATTGAGGGAGGGAAGGTAAATATTGACGGAAAT | 55 |

### **Table S9.** DNA template^7^, primers, target and the corresponding detector ssDNA for SARS-CoV-2 RNA detection.

| **Name** | **Sequence (5’-3’)** | **Len.** |
| --- | --- | --- |
| DNA template | TGGGGTTTTACAGGTAACCTACAAAGCAACCATGATCTGTATTGTCAAGTCCATGGTAATGCACATGTAGCTAGTTGTGATGCAATCATGACTAGGTGTCTAGCTGTCCACGAGTGCTTTGTTAAGCGTGTT | 132 |
| SARS-CoV-2 RNA fragment | UGGGGUUUUACAGGUAACCUACAAAGCAACCAUGAUCUGUAUUGUCAAGUCCAUGGUAAUGCACAUGUAGCUAGUUGUGAUGCAAUCAUGACUAGGUGUCUAGCUGUCCACGAGUGCUUUGUUAAGCGUGUU | 132 |
| Forward primer | **AATTCTAATACGACTCACTATAGGGAGAAGG**TGGGGTTTTACRGGTAACCT | 55 |
| Reverse primer | AACACGCTTAACAAAGCACTC | 30 |
| Target | CCATGATCTGTATTGTCAAGTCCATGGTAA |  |
| v4-COVID19 40-15 | ACCGTTGTAGCAATACTTCTTTGATTAGTAATAACATCAC**TTACCATGGACTTGA** | 55 |
| V8-COVID19 15-40 | **CAATACAGATCATGG**TCAACCGATTGAGGGAGGGAAGGTAAATATTGACGGAAAT | 55 |

### **Table S10. Backbone and basic variable oligos for the construction of nanoswitches and other oligos.**

| **Backbone oliogs** | | |
| --- | --- | --- |
| **#** | **Sequence (5’-3’)** | **Length** |
| 1 | AGAGCATAAAGCTAAATCGGTTGTACCAAAAACATTATGACCCTGTAATACTTTTGCGGG | 60 |
| 2 | AGAAGCCTTTATTTCAACGCAAGGATAAAAATTTTTAGAACCCTCATATATTTTAAATGC | 60 |
| 3 | AATGCCTGAGTAATGTGTAGGTAAAGATTCAAAAGGGTGAGAAAGGCCGGAGACAGTCAA | 60 |
| 4 | ATCACCATCAATATGATATTCAACCGTTCTAGCTGATAAATTAATGCCGGAGAGGGTAGC | 60 |
| 5 | TATTTTTGAGAGATCTACAAAGGCTATCAGGTCATTGCCTGAGAGTCTGGAGCAAACAAG | 60 |
| 6 | AGAATCGATGAACGGTAATCGTAAAACTAGCATGTCAATCATATGTACCCCGGTTGATAA | 60 |
| 7 | TCAGAAAAGCCCCAAAAACAGGAAGATTGTATAAGCAAATATTTAAATTGTAAACGTTAA | 60 |
| 8 | TATTTTGTTAAAATTCGCATTAAATTTTTGTTAAATCAGCTCATTTTTTAACCAATAGGA | 60 |
| 9 | ACGCCATCAAAAATAATTCGCGTCTGGCCTTCCTGTAGCCAGCTTTCATCAACATTAAAT | 60 |
| 10 | GGATAGGTCACGTTGGTGTAGATGGGCGCATCGTAACCGTGCATCTGCCAGTTTGAGGGG | 60 |
| 11 | ACGACGACAGTATCGGCCTCAGGAAGATCGCACTCCAGCCAGCTTTCCGGCACCGCTTCT | 60 |
| 12 | GGTGCCGGAAACCAGGCAAAGCGCCATTCGCCATTCAGGCTGCGCAACTGTTGGGAAGGG | 60 |
| 13 | CGATCGGTGCGGGCCTCTTCGCTATTACGCCAGCTGGCGAAAGGGGGATGTGCTGCAAGG | 60 |
| 14 | CGATTAAGTTGGGTAACGCCAGGGTTTTCCCAGTCACGACGTTGTAAAACGACGGCCAGT | 60 |
| 15 | GCCAAGCTTGCATGCCTGCAGGTCGACTCTAGAGGATCCCCGGGTACCGAGCTCGAATTC | 60 |
| 16 | GTAATCATGGTCATAGCTGTTTCCTGTGTGAAATTGTTATCCGCTCACAATTCCACACAA | 60 |
| 17 | CATACGAGCCGGAAGCATAAAGTGTAAAGCCTGGGGTGCCTAATGAGTGAGCTAACTCAC | 60 |
| 18 | ATTAATTGCGTTGCGCTCACTGCCCGCTTTCCAGTCGGGAAACCTGTCGTGCCAGCTGCA | 60 |
| 19 | TTAATGAATCGGCCAACGCGCGGGGAGAGGCGGTTTGCGTATTGGGCGCCAGGGTGGTTT | 60 |
| 20 | GTTGCAGCAAGCGGTCCACGCTGGTTTGCCCCAGCAGGCGAAAATCCTGTTTGATGGTGG | 60 |
| 21 | TTCCGAAATCGGCAAAATCCCTTATAAATCAAAAGAATAGCCCGAGATAGGGTTGAGTGT | 60 |
| 22 | TGTTCCAGTTTGGAACAAGAGTCCACTATTAAAGAACGTGGACTCCAACGTCAAAGGGCG | 60 |
| 23 | AAAAACCGTCTATCAGGGCGATGGCCCACTACGTGAACCATCACCCAAATCAAGTTTTTT | 60 |
| 24 | GGGGTCGAGGTGCCGTAAAGCACTAAATCGGAACCCTAAAGGGAGCCCCCGATTTAGAGC | 60 |
| 25 | TTGACGGGGAAAGCCGGCGAACGTGGCGAGAAAGGAAGGGAAGAAAGCGAAAGGAGCGGG | 60 |
| 26 | CGCTAGGGCGCTGGCAAGTGTAGCGGTCACGCTGCGCGTAACCACCACACCCGCCGCGCT | 60 |
| 27 | TAATGCGCCGCTACAGGGCGCGTACTATGGTTGCTTTGACGAGCACGTATAACGTGCTTT | 60 |
| 28 | CCTCGTTAGAATCAGAGCGGGAGCTAAACAGGAGGCCGATTAAAGGGATTTTAGACAGGA | 60 |
| 29 | ACGGTACGCCAGAATCCTGAGAAGTGTTTTTATAATCAGTGAGGCCACCGAGTAAAAGAG | 60 |
| 30 | TTGCCTGAGTAGAAGAACTCAAACTATCGGCCTTGCTGGTAATATCCAGAACAATATTAC | 60 |
| 31 | CGCCAGCCATTGCAACAGGAAAAACGCTCATGGAAATACCTACATTTTGACGCTCAATCG | 60 |
| 32 | TCTGAAATGGATTATTTACATTGGCAGATTCACCAGTCACACGACCAGTAATAAAAGGGA | 60 |
| 33 | CATTCTGGCCAACAGAGATAGAACCCTTCTGACCTGAAAGCGTAAGAATACGTGGCACAG | 60 |
| 34 | ACAATATTTTTGAATGGCTATTAGTCTTTAATGCGCGAACTGATAGCCCTAAAACATCGC | 60 |
| 35 | CATTAAAAATACCGAACGAACCACCAGCAGAAGATAAAACAGAGGTGAGGCGGTCAGTAT | 60 |
| 36 | TAACACCGCCTGCAACAGTGCCACGCTGAGAGCCAGCAGCAAATGAAAAATCTAAAGCAT | 60 |
| 37 | CACCTTGCTGAACCTCAAATATCAAACCCTCAATCAATATCTGGTCAGTTGGCAAATCAA | 60 |
| 38 | CAGTTGAAAGGAATTGAGGAAGGTTATCTAAAATATCTTTAGGAGCACTAACAACTAATA | 60 |
| 39 | GATTAGAGCCGTCAATAGATAATACATTTGAGGATTTAGAAGTATTAGACTTTACAAACA | 60 |
| 40 | CATTATCATTTTGCGGAACAAAGAAACCACCAGAAGGAGCGGAATTATCATCATATTCCT | 60 |
| 41 | GATTATCAGATGATGGCAATTCATCAATATAATCCTGATTGTTTGGATTATACTTCTGAA | 60 |
| 42 | TAATGGAAGGGTTAGAACCTACCATATCAAAATTATTTGCACGTAAAACAGAAATAAAGA | 60 |
| 43 | AATTGCGTAGATTTTCAGGTTTAACGTCAGATGAATATACAGTAACAGTACCTTTTACAT | 60 |
| 44 | CGGGAGAAACAATAACGGATTCGCCTGATTGCTTTGAATACCAAGTTACAAAATCGCGCA | 60 |
| 45 | GAGGCGAATTATTCATTTCAATTACCTGAGCAAAAGAAGATGATGAAACAAACATCAAGA | 60 |
| 46 | AAACAAAATTAATTACATTTAACAATTTCATTTGAATTACCTTTTTTAATGGAAACAGTA | 60 |
| 47 | CATAAATCAATATATGTGAGTGAATAACCTTGCTTCTGTAAATCGTCGCTATTAATTAAT | 60 |
| 48 | TTTCCCTTAGAATCCTTGAAAACATAGCGATAGCTTAGATTAAGACGCTGAGAAGAGTCA | 60 |
| 49 | ATAGTGAATTTATCAAAATCATAGGTCTGAGAGACTACCTTTTTAACCTCCGGCTTAGGT | 60 |
| 50 | GAAAACTTTTTCAAATATATTTTAGTTAATTTCATCTTCTGACCTAAATTTAATGGTTTG | 60 |
| 51 | AAATACCGACCGTGTGATAAATAAGGCGTTAAATAAGAATAAACACCGGAATCATAATTA | 60 |
| 52 | CTAGAAAAAGCCTGTTTAGTATCATATGCGTTATACAAATTCTTACCAGTATAAAGCCAA | 60 |
| 53 | CGCTCAACAGTAGGGCTTAATTGAGAATCGCCATATTTAACAACGCCAACATGTAATTTA | 60 |
| 54 | GGCAGAGGCATTTTCGAGCCAGTAATAAGAGAATATAAAGTACCGACAAAAGGTAAAGTA | 60 |
| 55 | ATTCTGTCCAGACGACGACAATAAACAACATGTTCAGCTAATGCAGAACGCGCCTGTTTA | 60 |
| 56 | TCAACAATAGATAAGTCCTGAACAAGAAAAATAATATCCCATCCTAATTTACGAGCATGT | 60 |
| 57 | AGAAACCAATCAATAATCGGCTGTCTTTCCTTATCATTCCAAGAACGGGTATTAAACCAA | 60 |
| 58 | GTACCGCACTCATCGAGAACAAGCAAGCCGTTTTTATTTTCATCGTAGGAATCATTACCG | 60 |
| 59 | CGCCCAATAGCAAGCAAATCAGATATAGAAGGCTTATCCGGTATTCTAAGAACGCGAGGC | 60 |
| 60 | ATTTTGCACCCAGCTACAATTTTATCCTGAATCTTACCAACGCTAACGAGCGTCTTTCCA | 60 |
| 61 | GAGCCTAATTTGCCAGTTACAAAATAAACAGCCATATTATTTATCCCAATCCAAATAAGA | 60 |
| 62 | AACGATTTTTTGTTTAACGTCAAAAATGAAAATAGCAGCCTTTACAGAGAGAATAACATA | 60 |
| 63 | AAAACAGGGAAGCGCATTAGACGGGAGAATTAACTGAACACCCTGAACAAAGTCAGAGGG | 60 |
| 64 | TAATTGAGCGCTAATATCAGAGAGATAACCCACAAGAATTGAGTTAAGCCCAATAATAAG | 60 |
| 65 | AGCAAGAAACAATGAAATAGCAATAGCTATCTTACCGAAGCCCTTTTTAAGAAAAGTAAG | 60 |
| 66 | CAGATAGCCGAACAAAGTTACCAGAAGGAAACCGAGGAAACGCAATAATAACGGAATACC | 60 |
| 67 | CAAAAGAACTGGCATGATTAAGACTCCTTATTACGCAGTATGTTAGCAAACGTAGAAAAT | 60 |
| 68 | ACATACATAAAGGTGGCAACATATAAAAGAAACGCAAAGACACCACGGAATAAGTTTATT | 60 |
| 69 | TTGTCACAATCAATAGAAAATTCATATGGTTTACCAGCGCCAAAGACAAAAGGGCGACAT | 60 |
| 70 | TCACCGTCACCGACTTGAGCCATTTGGGAATTAGAGCCAGCAAAATCACCAGTAGCACCA | 60 |
| 71 | TTACCATTAGCAAGGCCGGAAACGTCACCAATGAAACCATCGATAGCAGCACCGTAATCA | 60 |
| 72 | GTAGCGACAGAATCAAGTTTGCCTTTAGCGTCAGACTGTAGCGCGTTTTCATCGGCATTT | 60 |
| 73 | TCGGTCATAGCCCCCTTATTAGCGTTTGCCATCTTTTCATAATCAAAATCACCGGAACCA | 60 |
| 74 | GAGCCACCACCGGAACCGCCTCCCTCAGAGCCGCCACCCTCAGAACCGCCACCCTCAGAG | 60 |
| 75 | CCACCACCCTCAGAGCCGCCACCAGAACCACCACCAGAGCCGCCGCCAGCATTGACAGGA | 60 |
| 76 | GGTTGAGGCAGGTCAGACGATTGGCCTTGATATTCACAAACAAATAAATCCTCATTAAAG | 60 |
| 77 | CCAGAATGGAAAGCGCAGTCTCTGAATTTACCGTTCCAGTAAGCGTCATACATGGCTTTT | 60 |
| 78 | GATGATACAGGAGTGTACTGGTAATAAGTTTTAACGGGGTCAGTGCCTTGAGTAACAGTG | 60 |
| 79 | CCCGTATAAACAGTTAATGCCCCCTGCCTATTTCGGAACCTATTATTCTGAAACATGAAA | 60 |
| 80 | CCAGGCGGATAAGTGCCGTCGAGAGGGTTGATATAAGTATAGCCCGGAATAGGTGTATCA | 60 |
| 81 | CCGTACTCAGGAGGTTTAGTACCGCCACCCTCAGAACCGCCACCCTCAGAACCGCCACCC | 60 |
| 82 | TCAGAGCCACCACCCTCATTTTCAGGGATAGCAAGCCCAATAGGAACCCATGTACCGTAA | 60 |
| 83 | CACTGAGTTTCGTCACCAGTACAAACTACAACGCCTGTAGCATTCCACAGACAGCCCTCA | 60 |
| 84 | TAGTTAGCGTAACGATCTAAAGTTTTGTCGTCTTTCCAGACGTTAGTAAATGAATTTTCT | 60 |
| 85 | GTATGGGATTTTGCTAAACAACTTTCAACAGTTTCAGCGGAGTGAGAATAGAAAGGAACA | 60 |
| 86 | ACTAAAGGAATTGCGAATAATAATTTTTTCACGTTGAAAATCTCCAAAAAAAAGGCTCCA | 60 |
| 87 | AAAGGAGCCTTTAATTGTATCGGTTTATCAGCTTGCTTTCGAGGTGAATTTCTTAAACAG | 60 |
| 88 | CTTGATACCGATAGTTGCGCCGACAATGACAACAACCATCGCCCACGCATAACCGATATA | 60 |
| 89 | TTCGGTCGCTGAGGCTTGCAGGGAGTTAAAGGCCGCTTTTGCGGGATCGTCACCCTCAGC | 60 |
| 90 | CTTTTTCATGAGGAAGTTTCCATTAAACGGGTAAAATACGTAATGCCACTACGAAGGCAC | 60 |
| 91 | CAACCTAAAACGAAAGAGGCAAAAGAATACACTAAAACACTCATCTTTGACCCCCAGCGA | 60 |
| 92 | TTATACCAAGCGCGAAACAAAGTACAACGGAGATTTGTATCATCGCCTGATAAATTGTGT | 60 |
| 93 | CGAAATCCGCGACCTGCTCCATGTTACTTAGCCGGAACGAGGCGCAGACGGTCAATCATA | 60 |
| 94 | AGGGAACCGAACTGACCAACTTTGAAAGAGGACAGATGAACGGTGTACAGACCAGGCGCA | 60 |
| 95 | TAGGCTGGCTGACCTTCATCAAGAGTAATCTTGACAAGAACCGGATATTCATTACCCAAA | 60 |
| 96 | TCAACGTAACAAAGCTGCTCATTCAGTGAATAAGGCTTGCCCTGACGAGAAACACCAGAA | 60 |
| 97 | CGAGTAGTAAATTGGGCTTGAGATGGTTTAATTTCAACTTTAATCATTGTGAATTACCTT | 60 |
| 98 | ATGCGATTTTAAGAACTGGCTCATTATACCAGTCAGGACGTTGGGAAGAAAAATCTACGT | 60 |
| 99 | TAATAAAACGAACTAACGGAACAACATTATTACAGGTAGAAAGATTCATCAGTTGAGATT | 60 |
| 100 | TAAGAGCAACACTATCATAACCCTCGTTTACCAGACGACGATAAAAACCAAAATAGCGAG | 60 |
| 101 | AGGCTTTTGCAAAAGAAGTTTTGCCAGAGGGGGTAATAGTAAAATGTTTAGACTGGATAG | 60 |
| 102 | CGTCCAATACTGCGGAATCGTCATAAATATTCATTGAATCCCCCTCAAATGCTTTAAACA | 60 |
| 103 | GTTCAGAAAACGAGAATGACCATAAATCAAAAATCAGGTCTTTACCCTGACTATTATAGT | 60 |
| 104 | CAGAAGCAAAGCGGATTGCATCAAAAAGATTAAGAGGAAGCCCGAAAGACTTCAAATATC | 60 |
| 105 | GCGTTTTAATTCGAGCTTCAAAGCGAACCAGACCGGAAGCAAACTCCAACAGGTCAGGAT | 60 |
| 106 | TAGAGAGTACCTTTAATTGCTCCTTTTGATAAGAGGTCATTTTTGCGGATGGCTTAGAGC | 60 |
| 107 | TTAATTGCTGAATATAATGCTGTAGCTCAACATGTTTTAAATATGCAACTAAAGTACGGT | 60 |
| 108 | GTCTGGAAGTTTCATTCCATATAACAGTTGATTCCCAATTCTGCGAACGAGTAGATTTAG | 60 |
| 109 | TTTGACCATTAGATACATTTCGCAAATGGTCAATAACCTGTTTAGCTAT | 49 |

| **Variable oligos** | | |
| --- | --- | --- |
| **Name** | **Sequence (5’-3’)** | **Length** |
| Var 1 | AACATCCAATAAATCATACAGGCAAGGCAAAGAATTAGCAAAATTAAGCAATAAAGCCTC | 60 |
| Var 2 | GTGAGCGAGTAACAACCCGTCGGATTCTCCGTGGGAACAAACGGCGGATTGACCGTAATG | 60 |
| Var 3 | TTCTTTTCACCAGTGAGACGGGCAACAGCTGATTGCCCTTCACCGCCTGGCCCTGAGAGA | 60 |
| **Var 4** | TCTGTCCATCACGCAAATTA**ACCGTTGTAGCAATACTTCTTTGATTAGTAATAACATCAC** | **60** |
| Var 5 | **ATTCGACAACTCGTATTAAATCCTTTGCCCGAACGTTATT**AATTTTAAAAGTTTGAGTAA | 60 |
| **Var 6** | **TGGGTTATATAACTATATGTAAATGCTGATGCAAATCCAA**TCGCAAGACAAAGAACGCGA | 60 |
| **Var 7** | **GTTTTAGCGAACCTCCCGACTTGCGGGAGGTTTTGAAGCC**TTAAATCAAGATTAGTTGCT | 60 |
| **Var 8** | **TCAACCGATTGAGGGAGGGAAGGTAAATATTGACGGAAAT**TATTCATTAAAGGTGAATTA | **60** |
| Var 9 | GTATTAAGAGGCTGAGACTCCTCAAGAGAAGGATTAGGATTAGCGGGGTTTTGCTCAGTA | 60 |
| Var 10 | AGCGAAAGACAGCATCGGAACGAGGGTAGCAACGGCTACAGAGGCTTTGAGGACTAAAGA | 60 |
| Var 11 | TAGGAATACCACATTCAACTAATGCAGATACATAACGCCAAAAGGAATTACGAGGCATAG | 60 |
| Var 12 | ATTTTCATTTGGGGCGCGAGCTGAAAAGGTGGCATCAATTCTACTAATAGTAGTAGCATT | 60 |

| **Filler oligos** | | |
| --- | --- | --- |
| **Name** | **Sequence (5’-3’)** | **Length** |
| Var 4 filler | TCTGTCCATCACGCAAATTA | 20 |
| Var 5 filler | AATTTTAAAAGTTTGAGTAA | 20 |
| Var 6 filler | TCGCAAGACAAAGAACGCGA | 20 |
| Var 7 filler | TCGCAAGACAAAGAACGCGA | 20 |
| Var 8 filler | TATTCATTAAAGGTGAATTA | 20 |
| Var 9 filler | TAGCGGGGTTTTGCTCAGTA | 20 |

| **Other oligos** | |  |
| --- | --- | --- |
| Blocking | TCTCATGGCCCTTC | 14 |
| BtsCI cut site oligo | CTACTAATAGTAGTAGCATTAACATCCAATAAATCATACA | 40 |
